# Acetylation mimetic and null mutations within the filament core of P301L tau have varied effects on susceptibility to seeding and aggregation

**DOI:** 10.1101/2024.04.12.589253

**Authors:** Ethan D Smith, Jose Torrellas, Robert McKenna, Quan Vo, Mario Mietzsch, David R Borchelt, Stefan Prokop, Paramita Chakrabarty

## Abstract

The two most prominent post-translational modifications of pathologic tau are Ser/Thr/Tyr phosphorylation and Lys acetylation. Whether acetylation impacts the susceptibility of tau to templated seeding in diseases like Alzheimer’s disease (AD) and Progressive Supranuclear Palsy (PSP) is largely uncharacterized. Towards this, we examined how acetylation mimicking or nullifying mutations on five sites of tau (K311, K353, K369, K370, K375), located within the tau filament core, influenced the susceptibility of P301L (PL) tau to seeds from AD (AD-tau) or PSP (PSP-tau) in HEK293T cells. Acetyl-mimicking substitutions of individual Lys sites to Glutamine as well as mutation of all 5 sites together (PL+5K-Q and PL+5K-R) had inconsistent effects on tau seeding by AD-tau or PSP-tau seeds. Unexpectedly, mutating all 5 sites to Alanine (PL+5K-A) resulted in a tau variant that spontaneously aggregated. These aggregates were amorphous and yet able to propagate to naïve cells expressing P301L tau but not wild-type tau. Previously, we reported that a phospho-mimetic S305E mutation in PL tau abrogated seeding by AD-tau but not PSP-tau seeds in the HEK293T cells. To assess how changes in acetylation and phosphorylation together could influence seeding, we combined the S305E and the 5K-Q mutations in P301L tau, creating a variant that retained specificity for PSP-tau seeds over AD-tau seeds. Our findings indicate that phosphorylation of tau at Ser305 is a strong determinant of disease-specific tau templating, even in the presence of hyperacetylation within the fibril core domain. Overall, our findings suggest that acetylation could be a secondary modifier of misfolded hyperphosphorylated tau seeds.

## 1. Introduction

Tau is a microtubule binding and stabilizing protein that regulates axonal transport as well as participates in neuroplasticity (Wang & Mandelkow 2016; Koller & Chakrabarty). Aggregated tau is a characteristic hallmark of a broad category of neurodegenerative dementias, known as tauopathies, some examples of which are Alzheimer’s disease (AD), progressive supranuclear palsy (PSP), corticobasal degeneration (CBD), and primary age-related tauopathies (Gotz *et al*. 2019). Aggregated tau pathology in the brain is directly correlated with decline in mental acuity and extent of neurodegeneration, underpinning its pathological role in tauopathies and necessitating therapeutic targeting of pathological species of tau (Zhang *et al*. 2022).

Neuropathological and biochemical analysis of tau isolated from tauopathy patients show that it undergoes extensive post-translational modifications (PTMs) (Alquezar *et al*. 2020). Both tau that remains detergent-soluble as well as detergent-insoluble tau within neurofibrillary tangle (NFT) are extensively modified by PTMs, such as phosphorylation, acetylation, methylation, ubiquitination and glycosylation (Wesseling *et al*. 2020; Arakhamia *et al*. 2021; Kyalu Ngoie Zola *et al*. 2023). While phosphorylation is the major PTM that has been extensively studied, recently acetylation of Lys (K) residues has been revealed to mechanistically modify tau function and structure (Min *et al*. 2010; Cook *et al*. 2014a; Min *et al*. 2015; Tseng *et al*. 2021; Shin *et al*. 2021; Chakraborty *et al*. 2023). There are 44 lysine residues on tau that can be potentially acetylated. Previously, acetylation of residues K174 and K280/281 have been shown experimentally to induce tau aggregation (Min *et al*. 2015; Cohen *et al*. 2011; Tseng *et al*. 2021; Irwin *et al*. 2012), albeit through different mechanisms. These sites are found to be robustly acetylated in multiple types of tauopathies (Irwin *et al*. 2013; Min *et al*. 2015). However, acetylation or acetylation mimetics on many residues prone to acetylation (K259, K280, K290, K298, K321, K353) can also suppress tau fibrillization and pathology (Cook *et al*. 2014a; Chakraborty *et al*. 2023; Ajit *et al*. 2019). Acetylation could also prevent the availability of the Lys to other PTMs, such as methylation, glycation, ubiquitination and SUMOylation (Yang & Seto 2008), thus altering the structure-function profile of tau (Thomas *et al*. 2012). In addition, specific pseudo-acetylation of K321 or K280 has been shown to reduce tau hyperphosphorylation (Carlomagno *et al*. 2017; Ajit *et al*. 2019). Thus, acetylation could not only be associated with tau aggregation but also influence its PTM profile. Overall, there seems to be a complex series of biological outcomes when tau is acetylated leading to changes in ‘pathological’ phosphorylation patterns that are found in tau aggregates accumulating in neurodegenerative dementias.

In a previous study, we determined that the introduction of a phosphorylation mimetic mutation in Ser305 in P301L(PL)-tau modulated its ability to be seeded by tau aggregates derived from AD and PSP brains (Smith *et al*. 2023). Specifically, we found that S305E/PL tau was resistant to seeding by AD-tau seeds as compared to PSP-tau seeds. As disease-associated tau is post-translationally modified by acetylation (Alquezar *et al*. 2020), in this study, we examined whether acetylation on select sites would be crucial determinants of tau seeding. Indeed, two tour de force reports on tau cryo-EM revealed that acetylation preceded beta-sheet stacking of tau core protofilaments (Arakhamia *et al*. 2020; Fitzpatrick *et al*. 2017). We selected five acetylation sites (K311, K353, K369, K370, K375) that are present within the filament core domain (Fitzpatrick *et al*. 2017) and identified in late-stage AD patients with high frequency (Wesseling *et al*. 2020). Based on these findings, we hypothesized that acetylation on these sites within the tau core domain would facilitate templating of brain-derived tau seeds. As 0N/4R tau was enriched in the insoluble tau fraction in AD patients, we constructed recombinant 0N/4R PL tau with acetyl-mimetics (K→Q) and acetyl-nullifying substitutions (K→R or K→A) on these five sites. Using our previously described HEK293T cell model (Smith *et al*. 2023), we examined the aggregation state of these variants after seeding with detergent-insoluble tau seeds derived from AD or PSP patient brains. Our results indicate that PL tau, pseudo-acetylated tau and acetylation-deficient PL tau are equally sensitive to templating by AD-tau or PSP-tau seeds. Overall, our data shows that acetylation in the fibril core domain had minimal impact on the susceptibility of PL tau to seeded templating by brain-derived tau seeds.

## 2. Methods

### 2.1. Generation of tau acetyl-variant constructs

Acetyl-variant tau were generated in the human 0N4R PL tau backbone under contract with Genscript, carrying hybrid chicken β-actin promoter, Woodchuck promoter regulatory element and bovine poly A sequence. Institutional approval for the use of recombinant DNAs was obtained from Environmental Health & Safety Office at the University of Florida (Permit # BIO6317).

### 2.2. Purification of insoluble Tau from AD and PSP brains.□

The study was conducted according to the guidelines of the Declaration of Helsinki and approved by the Institutional Review Board of the University of Florida (IRB201600067). Human brain tissues from two AD cases and two PSP cases from the UF Brain Bank were selected for this study (**Table S1**). All cases were diagnosed based on accepted neuropathology criteria. Purification of pathological, insoluble tau from the temporal cortex of AD and lentiform nucleus of PSP cases were done as previously described (Smith *et al*. 2023). Briefly, for the purification of AD-tau and PSP-tau, 50-100 mg of temporal cortical gray matter or lentiform nucleus was homogenized in nine volumes (v/w) of high-salt buffer (10 mm□Tris with 0.8□m□NaCl, pH7.4) with 0.1% sarkosyl and 10% sucrose added, and centrifuged at 10,000 ×□g□for 10 min at 4°C. Pellets were re-extracted twice using the same high-salt buffer and the supernatants from all three extractions were filtered and pooled. Additional sarkosyl was added to the pooled supernatants to reach 1% and the samples were nutated for 1 h at 37°C. The samples were centrifuged at 150,000 ×□g□for 60 min at 15°C and the resulting 1% sarkosyl-insoluble pellets containing pathological tau were resuspended in PBS. The resuspended sarkosyl-insoluble pellets were further purified by a brief sonication using a handheld probe (Qsonica), followed by centrifugation at 100,000 ×□g□for 30 min at 4°C. The pellets were resuspended in PBS at 1/2 to 1/5 of the pre-centrifugation volume, sonicated, and spun at 10,000 ×□g□for 30 min at 4°C to remove large debris. The final purified supernatants contained insoluble, pathological tau, and are identified as AD-tau and PSP-tau in subsequent experiments. The final fraction was analyzed by Western blotting and sandwich ELISA for tau (Invitrogen #KHB0041) as previously characterized in (Smith *et al*. 2023). The total protein and total tau levels in S3 fraction are as follows: AD-tau (Patient A): total protein: 479.1µg/ml, total tau: 1054.2pg/ml; AD-tau (Patient B): total protein: 446.6µg/ml, total tau: 484.8pg/ml; PSP-tau (Patient A): total protein: 1555.5µg/ml, total tau: 663.6pg/ml; PSP-tau (Patient B): total protein: 1144.5µg/ml, total tau: 685.4pg/ml.

### 2.3. Cell Culture and Transfection

HEK293T cells were maintained in Dulbecco’s modified Eagle’s medium (Invitrogen, Carlsbad, CA) supplemented with 10% fetal bovine serum (FBS) and 100 U/ml penicillin/100 μg/ml streptomycin at 37°C and 5% CO2. For transfections, cells were plated on 24-well polystyrene plate with 1 mL of media. Once cells reached ∼60% confluency, 0.4 μg of plasmid DNA expressing 0N4R tau was combined with Lipofectamine 3000 (Thermo Fisher Scientific). For experiments designed to standardize expression of PL tau vs PL+5K-Q tau, a total of 1.6 µg was used for transfection. DNA corresponding to tau constructs was used as indicated (0.4, 0.8. 1.2 or 1.6 µg) and the total DNA content was adjusted to 1.6 µg using GFP DNA. In all cases, DNA mixture was incubated at room temperature for 10-15 minutes before adding dropwise to the media in each well and placed in the incubator for 30 minutes. For seeding, 0.5 μg of detergent-insoluble preparations of human brain was incubated with Lipofectamine reagent for 20 minutes and added to the cells. Cells were harvested 48 hours after transfection and fractionated.

### 2.4. Biochemical Fractionation of HEK293T cells

Cells were harvested in 50 μL of High Salt Buffer (HSB: 50 mM Tris-HCl, pH 7.4, 250 mM NaCl, 2 mM EDTA, 1% Triton X-100, 20 mM NaF) and a cocktail of protease and phosphatase inhibitors (Pierce #A32959). Samples were sedimented at 150,000 x g for 30 minutes at 4°C and the supernatants collected, washed and sedimented again. Supernatants were removed and the pellets were resuspended in 50 μL of HSB. SDS sample buffer (final concentration of 250 mM Tris-Cl, pH 6.8, 5% β-Mercaptoethanol, 0.02% Orange G, 10% SDS, 30% glycerol) was added to the collected supernatants and resuspended pellets, referred to as the detergent-soluble and detergent-insoluble fractions respectively. To resuspend the pellet, detergent-insoluble samples were sonicated in HSB buffer and resuspended in 2x SDS-PAGE loading buffer. These samples were then heated at 90°C for 10 minutes.

The primary outcomes of the study are measured as Relative Tau Aggregation and Seeded AT8 Ratio. The relative tau aggregation was calculated as [(total tau in detergent insoluble fraction)/(total tau in detergent soluble fraction□+□total tau in detergent insoluble fraction)]*100. Seeded AT8 Ratio denotes normalized AT8 values, calculated as [AT8 signal in detergent-insoluble seeded fraction / AT8 signal in detergent-soluble seeded fraction] and expressed as percentage.

### 2.5. Preparation of Secondary Seeds and Passaging of 5K-A tau

Briefly, HEK293T cells transfected with PL+5K-A tau were seeded with 0.5 μg of detergent-insoluble preparations of human brain or left unseeded and harvested after 48 hours. Cellular material was then fractionated and the detergent-insoluble fraction was used for secondary passaging experiments. Detergent-insoluble seeds were then characterized via immunoblotting and ImmunoEM, followed by secondary passaging. The secondary seeds are labeled as PL+5KA-AD, PL+5KA-PSP and PL+5KA-U to denote that the seeds were derived from HEK293T cells expressing PL+5K-A and seeded with AD-tau or PSP-tau or unseeded. For secondary passaging experiments, HEK293T cells transfected with WT tau or PL tau were seeded with 50ng of these secondary seeds and harvested after 48 hours. Cellular material was further fractionated into detergent soluble and insoluble fractions, followed by characterization of seeded tau via immunoblot.

### 2.6. Tau co-sedimentation with paclitaxel-stabilized microtubules

HEK293T cells were lysed in 100 μl of PEM buffer (80 mM□PIPES, pH 6.8, 1 mM□EGTA, 1 mM□MgCl2) supplemented with 0.1% Triton X-100, 2 mM□GTP, 20 μM□paclitaxel, and a mix of protease inhibitors as described previously (Smith *et al*. 2023). For the “without paclitaxel” condition, cells were lysed in PEM buffer supplemented with 0.1% Triton X-100 and a mix of protease and phosphatase inhibitors. Cell lysates were incubated in a 37 °C water bath for 30 min and then centrifuged at 100,000 ×□g□for 30 min to pellet microtubules (MT). Supernatant was transferred to a new tube, and the pellet (MT fraction with bound proteins) was resuspended in PEM buffer. The pellet fraction was briefly sonicated, and SDS gel loading buffer was added to both fractions. Equivalent amounts of supernatant and pellet were loaded on SDS-polyacrylamide gels for Western blot analysis. Percent MT bound tau was calculated as pellet/(supernatant + pellet) × 100. Tubulin polymerization was calculated as a ratio of β-tubulin in the Pellet (P):Supernatant (S) fraction.

### 2.7. Western blotting

8 μl of detergent-soluble and 15 μl of detergent-insoluble samples were loaded on 4-20% Tris-Glycine gels (Invitrogen #XP04205) and separated for 1.5hrs at 100V. Blots were then transferred to 0.45 μm PVDF membranes (Millipore #IPFL85R) for 2hrs at 300 mAmps before being blocked in 0.5% Casein (Hammarsten grade; ThermoFisher #J12840.Q1) dissolved in TBS buffer for 1 hr. Blots were incubated in appropriate primary antibodies diluted in 0.5% casein in TBS (**Table S2**) overnight followed by at least 3 washes in 1x TBS buffer. IRDye-labeled secondary antibodies (IRDye® 800CW Goat anti-Mouse IgG Secondary Antibody or IRDye® 680RD Goat anti-Rabbit IgG Secondary Antibody, LICORbio) were mixed and diluted in 0.5% casein (1:20,000) with 0.005% SDS. Bots were incubated with this mix antibody for 1 h at room temperature. Membranes were washed three times in 1× TBS for 5 min, and protein bands were detected using the multiplex Li-Cor Odyssey Infrared ODY-2536 Imaging system (Li-Cor Biosciences, Lincoln, NE, USA).

### 2.8. Immuno-gold, negative-stain electron microscopy

Tau filaments from detergent-insoluble HEK293T cell lysates were deposited on 300 mesh carbon film-coated copper grids (Ted Pella 01843-F) for 10 minutes, blocked for 15□min with PBS□+□2% BSA, and incubated with anti-tau CP27 antibody (1:500) in PBS + 1% BSA overnight. Grids were washed with blocking buffer and incubated with 10□nm gold-conjugated anti-mouse IgG (Sigma G7652) diluted 1:50 in PBS + 1% BSA for 2 hrs. The grids were then washed with water and stained with 1% uranyl acetate for 60□s. Images were acquired using a Tecnai G2 Spirit at 120□kV (University of Florida ICBR Electron Microscopy Core Facility, RRID:SCR_019146)

### 2.9. In Silico Predictions

Bioinformatic modeling of tau acetyl-mimetics were carried out using the ChimeraX program (Meng *et al*. 2023). Using the previously published tau core domains for AD (V306-F378) and PSP (G272-N381) (Fitzpatrick *et al*. 2017; Shi *et al*. 2021), individual amino acid residues corresponding to PL+5K-Q were in silico substituted and the intramolecular environmental distances were measured against adjacent residues.

### 2.10. Statistical Analysis

Western blot signals were quantified as mean ± S.E.M. based on densitometric analysis using ImageJ. All analysis was using GraphPad software (GraphPad Prism version 10.5.0 for Windows, Boston, Massachusetts USA, www.graphpad.com). For statistical comparisons, we performed both one-way and two-way analysis of variance (ANOVA) with either a Dunnett’s or Sidak’s multiple comparisons test, to compare each group to the control. No blinding was done in this study.

## 3. Results

### 3.1. Pseudo-acetylated tau shows variable aggregation in presence of tau seeds

A proteomic analysis of detergent insoluble tau extracted from a well-characterized cohort of AD patients identified major phosphorylation and acetylation sites present on tau protein across various stages of the disease (Wesseling *et al*. 2020). This study showed that acetylation modifications present in AD patients were mostly clustered in the MTBR and C-terminus region of tau that overlaps with the AD-tau filament core domain. To understand how acetylation within the AD-tau filament core domain may affect prion-like propagation, we selected 5 of these lysine sites (K311, K353, K369, K370 and K375) for mutating to acetyl-mimicking amino acids. To test the hypothesis that hyperacetylation on these residues would positively reinforce formation of tau aggregates, we used a robust HEK293T cell seeding model we have previously described (Smith *et al*. 2023). Our aim was to determine whether AD-tau seeds and PSP-tau seeds extracted from human postmortem brains could successfully template on tau that contain acetyl-mimicking or acetyl-nullifying residues on these potential AcK sites.

To investigate potential patient heterogeneity in seeding, we used seed preparations from two individuals with AD and two individuals with PSP (referred to as Patients A and B for each diagnosis: AD-A, AD-B, PSP-A, PSP-B). All cases used for this study were neuropathologically confirmed and their seeding profiles were characterized in an earlier study (**Table S1**) (Smith *et al*. 2023). As wild type 0N4R tau is not templated efficiently in cellular models (Smith *et al*. 2023), we used human 0N/4R PL tau as the recipient tau with and without additional pseudo-acetylation mutations. Detergent-insoluble AD and PSP brain lysates (∼0.5 μg of detergent-insoluble protein) were used to seed HEK293T cells expressing each pseudoacetyl construct. After 2 days, cell lysates were fractionated, separated by SDS-PAGE, and probed with a total tau antibody (CP27) to reveal the amount of tau that became misfolded for each construct, represented by ‘Relative Tau Aggregation’ in the detergent-insoluble cell fraction. We also used AT8 (pSer202/Thr205) antibody because it is highly reactive to human neurofibrillary tangles (NFT) and is generally non-reactive in normal brain tissue (Mercken *et al*. 1992), calculating the relative phosphorylation levels of the detergent insoluble tau to soluble tau and expressed ‘Seeded AT8 Ratio’.

We used 0.5 μg of AD lysate from Patient A (**Fig. 1a-f**) or Patient B (**Fig. S1a-f**) to seed HEK293T cells expressing individual pseudo-acetyl mutations (K311, K353, K369, K370, and K375) on PL tau. After 2 days, cell lysates were fractionated, separated by SDS-PAGE, and probed with CP27 antibody and phosphorylation-specific AT8 antibody to reveal levels of insoluble tau. The individual pseudo-acetyl tau constructs expressed similarly and showed comparable AT8 levels in the soluble fraction (**Fig. 1a, d; Fig. S1a, d**). Although we found some variation in the efficacy of seeding these tau variants with seeds from AD Patient A (**Fig. 1c**), these differences were limited with seeds from AD Patient B (**Fig. S1c**). Specifically, for Patient A, we observed that PL+K369Q and PL+375Q had lower seeded tau compared to PL-tau (**Fig. 1c**; p<0.05). The levels of AT8 reactive insoluble tau were also variable but none of the AD-tau seeded pseudo-acetyl tau showed differential AT8 phosphorylation relative to each other or parent PL tau (**Fig. 1f; Suppl. Fig. S1f**).

**Figure 1:**
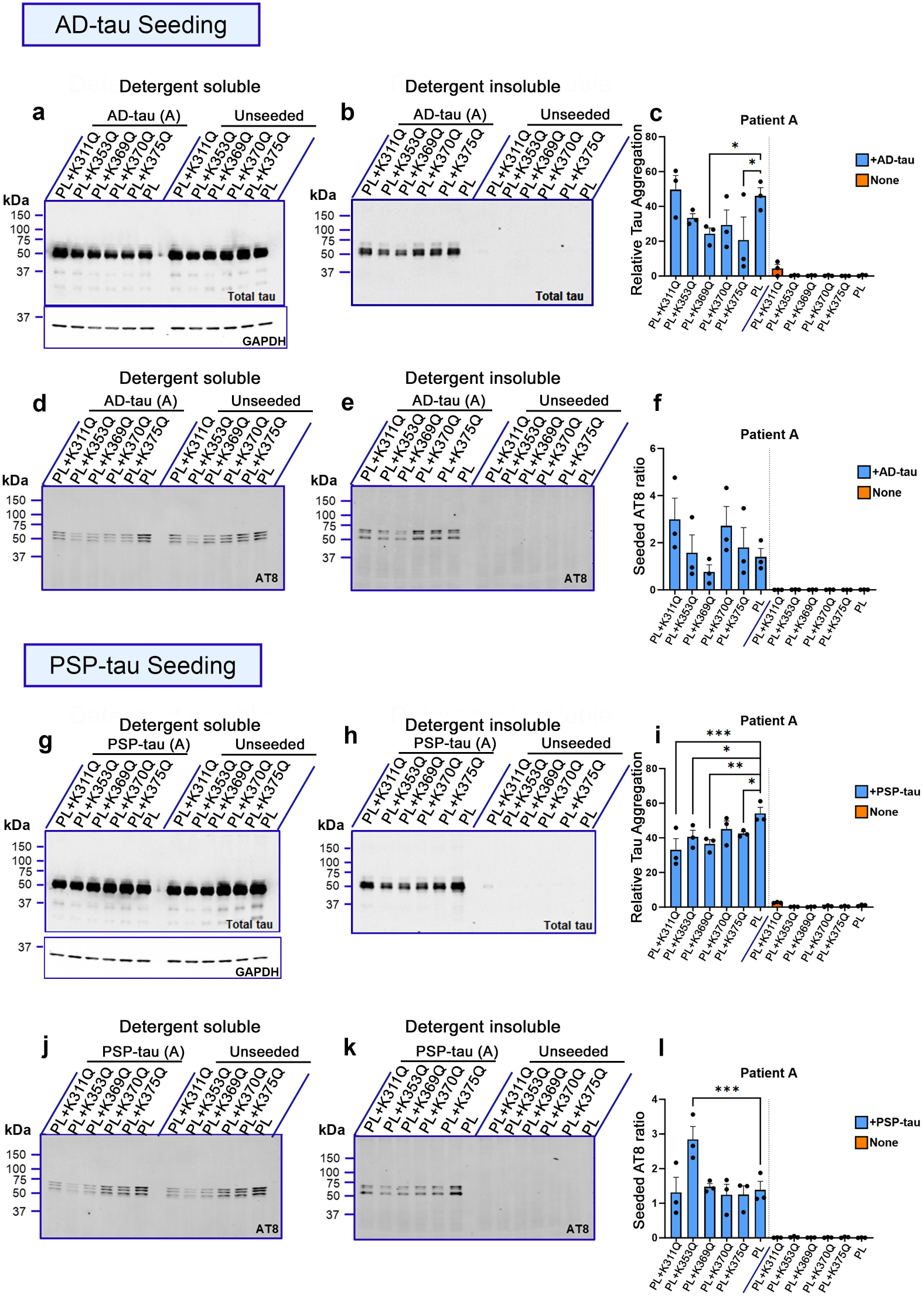
Individual pseudo-acetyl variants in the tau filament core domain show variable seeding propensity. a-c. HEK293T cell seeding assay using the acetyl variants on P301L (PL) tau backbone, seeded with AD-tau seeds or left unseeded. Samples were fractionated into detergent-soluble and detergent-insoluble lysates and probed for total tau (t-tau) (a, b). Quantification of relative tau aggregation (ratio of insoluble tau to soluble tau) for each tau variant is shown (c). GAPDH immunoblot indicates protein loading control in the detergent soluble fraction. d-f. Representative AT8 immunoblot of individual acetyl-mimetic PL tau seeded with AD:Patient A tau seeds or left unseeded (d, e). Seeded AT8 Ratio denotes AT8 levels in insoluble seeded fraction normalized to corresponding soluble seeded fraction for each variant (f). g-i. Representative total tau immunoblot of detergent-soluble or detergent-insoluble cell lysates of individual acetyl-mimetic PL tau seeded with PSP:Patient A tau seeds or left unseeded (g, h). Quantification of relative aggregation for each variant is shown (i). GAPDH immunoblot indicates protein loading control. j-l. Representative AT8 immunoblots (j, k) and AT8 Ratio (l) of individual acetyl-mimetic PL tau seeded with PSP:Patient A tau seeds. Relative molecular masses (kDa) are indicated on the left of each blot. All graphs represent mean ± S.E.M. N=3 for each experimental replicate. 1-way ANOVA with Sidak’s multiple comparisons test, with single pooled variance. *p<0.05, **p<0.01, ***p<0.001. PL, P301L tau; AD, Alzheimer’s disease; PSP, Progressive supranuclear Palsy.

We next tested how PSP-tau seeds from two different patients affected seeding efficiency of the individual pseudo-acetyl PL tau constructs (**Fig 1g-l**; **Fig. S1g-l**). We found inter-patient variability in seeding efficiency in this case also. Specifically, most all the individual acetyl variants (PL+K311Q – p<0.001, PL+K353Q – p<0.05, PL+K369Q – p<0.01, and PL+K375Q – p< 0.05) showed lower seeding efficiency with PSP-tau seeds from Patient A compared to PL tau (**Fig. 1g-i**; p<0.05). However, PSP Patient B seeds showed no differences in seeding on any of the acetyl tau variants compared to PL tau (**Fig. S1g-i**). The induction of AT8 reactivity was comparable for these acetyl variants following seeding in both cases, except for PL+K353Q tau that showed higher AT8 levels compared to PL tau following seeding in both cases (p<0.001 for Patient A; p<0.05for Patient B) (**Fig. 1l**; **Fig. S1l**). None of the unseeded pseudo-acetyl tau variants displayed detectable AT8 or total tau in the insoluble cellular fraction (**Fig. 1i, l; Suppl. Fig. S1i, l**). Overall, the individual pseudo-acetyl tau variants showed variable propensities for seeded aggregation, based on seed origins.

### 3.2. Combinatorial hyperacetyl-modified tau shows equivalent aggregation properties in the presence of tau seeds

Because tau is generally found in hyperacetylated form (Alquezar *et al*. 2020), we reasoned that if we combinatorially substitute all the AcK sites to Gln, we would be able to recapitulate seeding activity in brains that are well into the pathogenic cascade. In this study, all 5 lysine residues were mutated combinatorically to either K→Q (acetyl-mimicking or acetyl-plus, PL+5K-Q), K→R (acetyl-null, PL+5K-R), or K→A (acetyl-null, PL+5K-A) on the 0N/4R PL tau backbone to investigate changes in the propagation of brain derived seeds. Examining tau levels in the insoluble fraction following seeding from two different patients, we found that for both AD-tau and PSP-tau seeds, the cells expressing PL+5K-Q tau had dramatically lower insoluble tau relative to parent PL tau (p<0.0001) (**Fig. 2b, c, e, f**; **Fig. S2b, c, e, f**). However, the expression levels of the PL+5K-Q tau variant in the soluble fraction from both seeded and unseeded cells were somewhat lower than the parent PL tau (**Fig. 2a**), raising the concern that this variation by itself could explain the reduced seeding in PL+5K-Q. We confirmed that PL+5K-Q and PL-tau expressed at different levels with varying DNA concentrations (**Fig. S2g**), and then we repeated AD-tau seeding using progressively increasing DNA concentrations of PL and PL+5K-Q constructs (**S2h-l**). We observed that while the PL+5K-Q tau showed lower seeding efficiency relative to PL tau under limiting conditions of transfected DNA (0.4µg), this effect vanished when expression of PL tau and PL+5K-Q tau were increased (**Fig S2h-l**). In addition, we observed that relative tau aggregation did not increase linearly with higher transfected amounts of the substrate (**Fig. S2l**), suggesting that the seeds are limiting in this scenario. This finding suggests that the poor seeding of PL+5K-Q tau seen when 0.4 µg of DNA was used in transfection was essentially due to lower steady-state levels of protein rather than an inhibitory effect by the hyperacetylation mimetic mutations.

**Figure 2:**
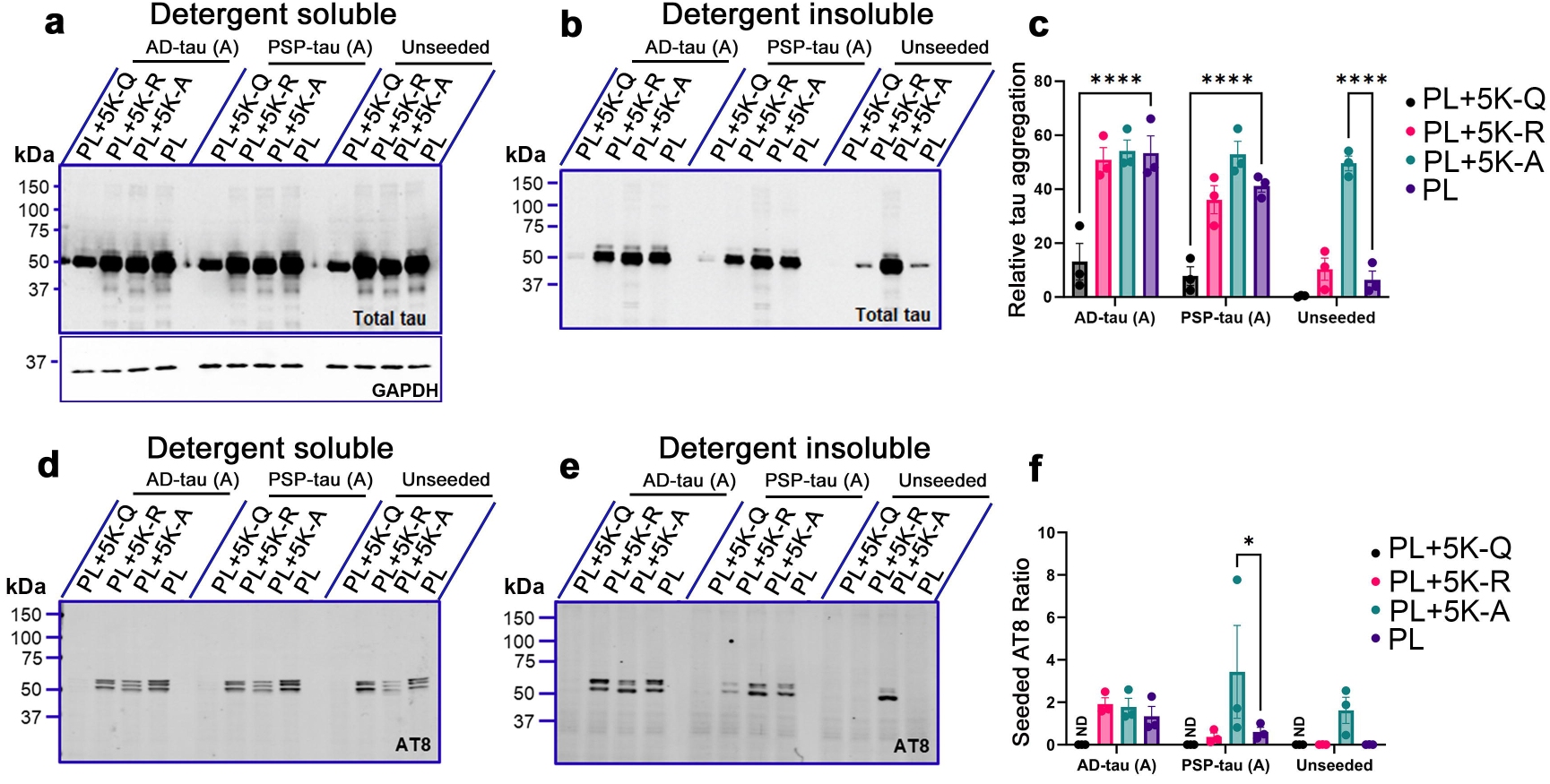
Comparative effect of combinatorial acetyl variants in the tau core domain on seed-induced PL tau aggregation. a-c. HEK293T cell seeding assay using the acetyl variants on P301L (PL) tau backbone, seeded with AD-tau seeds or PSP-tau seeds. Samples were fractionated into detergent-soluble and detergent-insoluble lysates and probed for total tau (t-tau) and p-tau (AT8). Representative immunoblot (a, b) and relative tau aggregation (ratio of insoluble tau to soluble tau) (c) of acetyl-tau variants seeded with AD:Patient A tau seeds or PSP:Patient A tau seeds or left unseeded. GAPDH immunoblot indicates protein loading control in the detergent soluble fraction. d-f. Representative immunoblots (d, e) of detergent-soluble and detergent-insoluble lysates and Seeded AT8 Ratio (f) of tau acetyl variants seeded with AD:Patient A tau seeds or PSP:Patient A tau seeds or left unseeded. Relative molecular masses (kDa) are indicated on the left of each blot. All graphs represent mean ± S.E.M. N=3 for each experimental replicate. ND, not done. 2-way ANOVA with Dunnett’s multiple comparisons test, with single pooled variance. *p<0.05, ****p<0.0001. N.D., not detected; PL, P301L tau; AD, Alzheimer’s disease; PSP, Progressive supranuclear Palsy.

When we examined the acetylation-null mimetic PL+5K-R tau under the same conditions of seeding, we observed similar levels of tau aggregation as well as seeded AT8 levels relative to PL tau **(Fig. 2b, c, e, f; Fig. S2c, f**). Although the PL+5K-A variant also showed induced aggregation levels similar to parental PL tau, we observed that PL+5K-A tau spontaneously formed AT8-positive insoluble inclusions in the absence of any brain-derived tau seeds (unseeded: **Fig. 2b, c, e, f**; **Fig. S2b, c, e, f**). AT8 levels were also increased in PSP-tau seeded PL+5K-A expressing cells relative to PL (**Fig. 2f**, p<0.05; **Fig. S2f**, p<0.01), though this was not noted in AD-tau seeded cells. Overall, our data indicates that substitution of these Lys residues with Ala renders tau prone to spontaneous aggregation.

### 3.3. PL tau with acetyl-null 5K-A substitutions in the tau filament core produce seeding competent amorphous aggregates

We showed that PL+5K-A tau has the propensity to form AT8-positive insoluble tau. Here, we have examined whether exposure of PL+5K-A tau to AD-tau or PSP-tau seeds produces a detectable strain variation. HEK293T cells expressing PL+5K-A tau were exposed to AD-tau seeds, PSP-tau seeds or left unseeded to generate aggregated tau (**Fig. 3a**; ‘primary passage’). The detergent-insoluble PL+5K-A tau aggregates from these cells were first analyzed by immunogold electron microscopy (immuno-EM) using anti-total tau CP27 antibody and secondary antibodies labeled with 10nm immunogold particles (**Fig. 3b, Fig. S3**). In all preparations, we detected similar types of amorphous deposits labeled by the gold particles, indicating that the presence of human brain-derived tau seeds did not alter the morphology of spontaneously-aggregating PL+5K-A (**Fig. 3b**). We confirmed that in the absence of primary antibody, there was no substantial immunogold labeling, indicating staining specificity (**Fig. S3d**).

**Figure 3:**
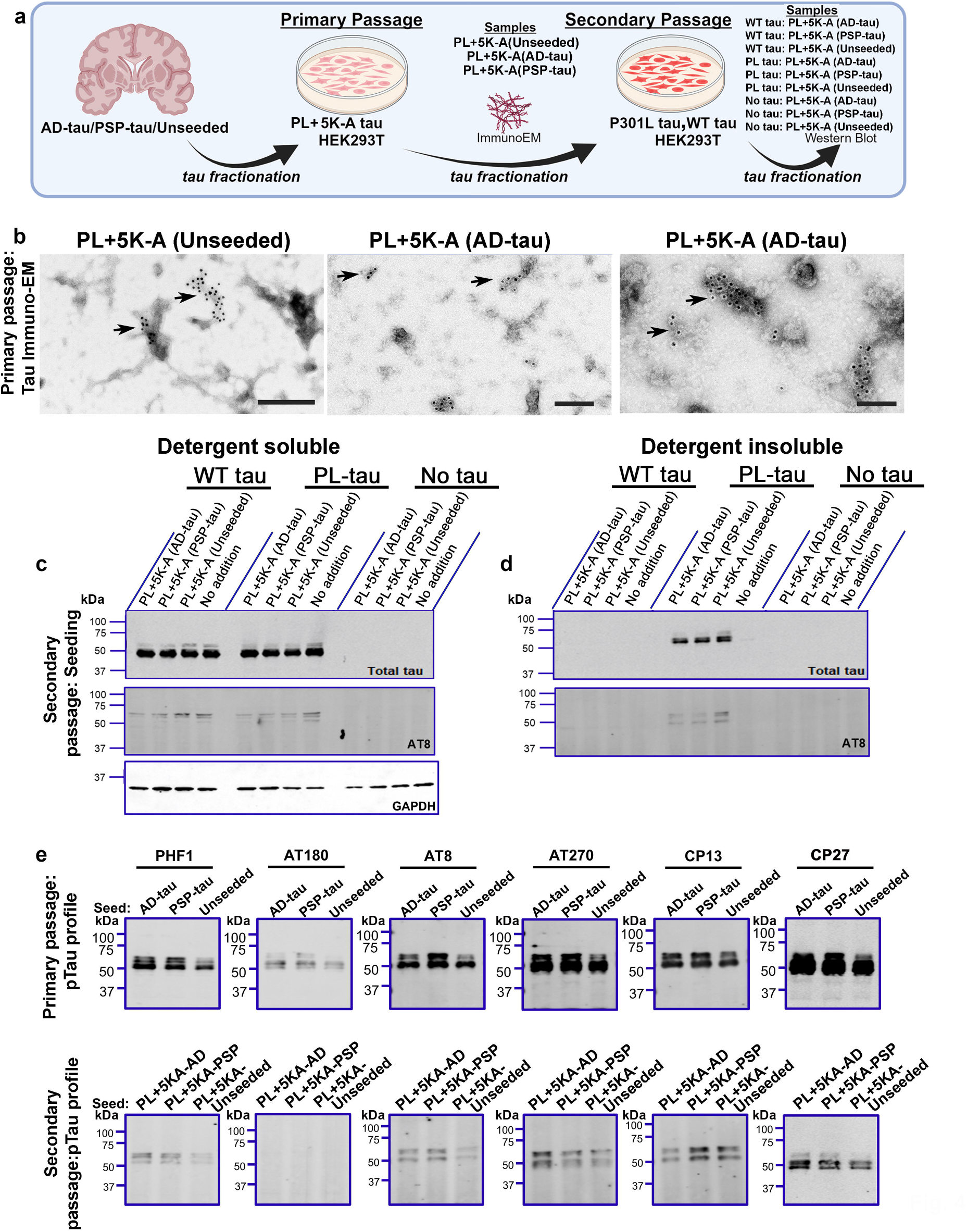
Acetyl nullifying P301L tau displays unique self-aggregating properties. a. Schematic description of secondary passaging paradigm. Briefly, HEK293T cells expressing PL+5K-A tau constructs were either seeded with AD-tau, PSP-tau, or left unseeded. The detergent-insoluble fraction was harvested and used again (‘secondary 5K-A seeds’) for seeding of WT tau, PL tau or No tau conditions. b. Representative immuno-EM images of total tau antibody-stained detergent-insoluble 5K-A tau seeded with either AD-tau, PSP-tau, or unseeded. The preparations were stained with 10nm gold conjugated secondary antibody followed by negative staining with 1% uranyl acetate. Arrows depict 10nm gold particles. Additional EM images for each condition are shown in **Fig. S3**. c-d. Representative total tau and AT8-tau blots depicting outcomes of secondary passaging of PL+5K-A seeds on HEK293T cells transfected with WT tau, PL tau or no DNA added (No tau). GAPDH immunoblot indicates protein loading control in the detergent soluble fraction. e. Characterization of the phosphorylation profile (using PHF1, AT180, AT8, AT270 and CP13 antibodies) for primary passaged AD-tau, PSP-tau or unseeded lysates on PL+5K-A (top row) which were then used for secondary passaging on PL tau (bottom row) with unseeded lane receiving no seeds. CP27 denotes total tau. Representative blots from n=2 replicates. WT, wild type; PL, P301L tau; AD, Alzheimer’s disease; PSP, Progressive supranuclear Palsy.

We then used the secondary passaging strategy commonly used in the prion field (Makarava *et al*. 2020) to assess if PL+5K-A tau had acquired the unique seeding properties of the original seeds that it was exposed to (**Fig. 3a**; ‘secondary passage’). The same preparations analyzed by EM were used as tau seeds on HEK293T cells expressing WT tau or PL tau (**Fig. 3c**). For these secondary passaging experiments, the PL+5K-A seeds (AD-tau primed, PSP-tau primed or naive) were diluted 1:10 (∼50ng of total protein), as the undiluted samples caused cells to lift off within 24 hours of application. We found that all forms of PL+5K-A seeds (AD-tau primed or PSP-tau primed or naive) were able to secondarily seed PL tau and induce modest levels of AT8 positivity (**Fig. 3d**). Cells expressing WT tau did not develop aggregates following exposure to these seeds under the same experimental conditions (**Fig. 3d**). We further examined the phosphorylation patterns of PL+5K-A seeds from primary passage seeding and secondary passaging (**Fig. 3e**). In the primary passage, detergent-insoluble PL+5K-A showed phosphorylation on epitopes defined by reactivity to PHF1, AT180, AT8, AT270, and CP13 antibodies, irrespective of whether PL+5K-A tau was seeded with AD-tau, PSP-tau or left unseeded (**Fig. 3e**, top panel). Notably, unseeded PL+5K-A showed lower levels of the 65kDa hyperphosphorylated tau than AD-tau and PSP-tau seeded 5K-A. In the secondary passaged seeds, the resulting detergent-insoluble tau showed some level of phosphorylation on PHF1, AT8, AT270, and CP13 epitopes. Notably, in all cases the secondary passage generated tau seeds completely lacked phosphorylation on the AT180 epitope (**Fig. 3e**, bottom panel). These findings suggest that PL+5K-A tau may be a unique strain that aggregates spontaneously at a high frequency independently of any exposure to seeding aggregates. Notably, this variant retains a high propensity for phosphorylation though our findings hint that lack of acetylation may modulate phosphorylation at AT180 (pThr 231) epitope.

### 3.4. Differential microtubule co-sedimentation patterns of acetyl tau variants

Tau plays a key role in assembly of tubulin protein and further in maintaining structural stability of axonal microtubules (Cook *et al*. 2014b). To investigate whether the acetylation of key residues within the tau filament core domain affects how it interacts with microtubules, we performed a sedimentation assay in the presence or absence of the microtubule-stabilizing compound Paclitaxel (**Fig. 4, Fig. S4**). HEK293T cells expressing pseudo-acetyl tau were harvested with high salt buffer supplemented with Paclitaxel and GTP, followed by ultracentrifugation to reveal free tau (S, soluble) or tau co-sedimenting with polymerized microtubules (P, pellet). Each of these fractions were probed for total tau and β-tubulin (β-tub) (**Fig. 4; Fig. S4**). In the absence of paclitaxel, the majority of tubulin is unpolymerized and thus remains soluble, whereas in the presence of paclitaxel, the majority of tubulin is polymerized and is precipitated in the pellet fraction. First, we confirmed that none of the combinatorial acetyl-tau variants affected the ability of paclitaxel to polymerize and precipitate the tubulin multimers when co-expressed (**Fig. S4a**). As expected, presence of paclitaxel resulted in higher proportion of β-tubulin in the pellet fraction, when compared to conditions without paclitaxel, indicating microtubule polymerization (**Fig. S4a**). Notably, the PL+5K-Q tau remained in the soluble fraction in paclitaxel plus condition, indicating that PL+5K-Q tau did not co-sediment with polymerized microtubules efficiently (p<0.05 compared to PL tau) (**Fig. 4a, b**). This suggests that the acetyl-mimic 5K-Q tau binds to polymerized tubulin poorly. On the other hand, PL+5K-R tau and PL tau showed similar propensity to co-sediment with tubulin in the pellet fraction in the presence of paclitaxel (**Fig. 4a, b**). Compared to PL tau, the levels of PL+5K-A tau were over-represented in the insoluble fraction in the paclitaxel plus condition (**Fig. 4a, b**). Because PL+5K-A was similarly over-represented in the pellet fraction in the absence of paclitaxel when the microtubules remained mostly unpolymerized and soluble (**Fig. 4c, d**: ∼35% in both paclitaxel conditions), this suggests that sedimentation of PL+5K-A tau occurs due to its inherent insolubility. Indeed, comparing the paclitaxel plus and minus conditions (**Fig. 4b** vs **Fig. 4d**), it can be assumed that PL+5K-A probably does not associate with microtubules efficiently. Neither GFP expression nor naïve HEK293T cells showed detectable tau co-sedimentation in this assay (**Fig. 4e-f**).

**Figure 4:**
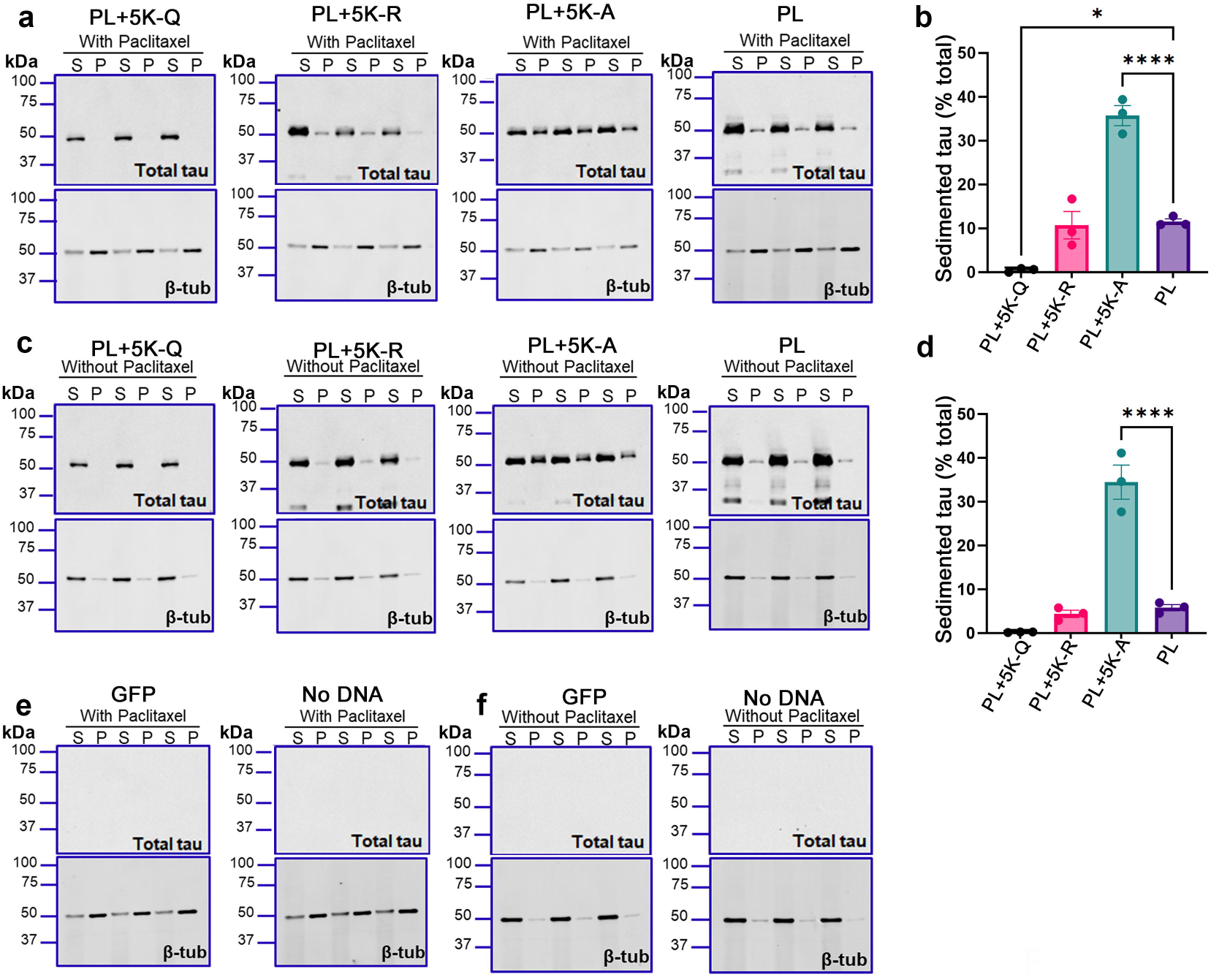
Comparative microtubule association properties of combinatorial acetyl-substituted tau. HEK293T cell-based microtubule binding assay was performed in the presence or absence of Paclitaxel with cells transfected to express different PL tau with acetyl-substituted epitopes. Antibody specific to β-tubulin was used to detect microtubule under the same conditions, while total tau antibody (CP27) was used to detect the amount of sedimented tau. a-b. Representative blots and quantification of sedimentation efficiency of PL+5K-Q, PL+5K-R and PL+5K-A compared to parent PL tau in paclitaxel-stabilized conditions. c-d. Representative blots and quantification of sedimentation of PL+5K-Q, PL+5K-R and PL+5K-A compared to parent PL tau under native conditions (no paclitaxel added). e-f. Representative blots of sedimented tau in the absence of tau expression, exemplified by presence of GFP or no DNA added to transfection (e, paclitaxel added and f, no paclitaxel). The relative molecular masses of protein markers are indicated on the left. All graphs represent mean ± S.E.M. N=3 for each experimental replicate. 1-way ANOVA with Dunnett’s multiple comparisons test, with single pooled variance. *p<0.05, ****p<0.0001. Blots were probed for total tau or β-tubulin (β-tub). PL, P301L tau; S, supernatant fraction; P, pellet fraction.

We also investigated the relative microtubule-binding affinities of the individual pseudo-acetyl variants (**Fig. S4b-e**). All of these pseudo-acetyl variants showed equivalent levels of sedimentation in the pellet fraction in the presence of paclitaxel (**Fig. S4b-c**), except for PL+K370Q which showed slightly increased presence in the pellet fractions (p=0.0271 compared to PL tau). In the absence of paclitaxel, we did not observe any of these individual pseudo-acetyl variants to be more prone to inherent insolubility, although the PL+K370Q variant had the highest level of sedimented tau (**Fig. S4d, e**). Together, these results indicate that neutralizing the positive charge in the P301L tau filament core reduces its propensity to associate with polymerized microtubules consistent with previous reports (Cohen *et al*. 2011), though individual acetyl modifications within the filament core domain has minimal effects on microtubule association.

### 3.5. Inclusion of phospho-mimicking Ser305Glu substitution enables hyperacetylated tau to show disease-specific seeding characteristics

Previous studies have indicated that hyperacetylation on specific residues is related to phosphorylation patterns and these together regulate tau polymerization (Carlomagno *et al*. 2017). This association prompted us to consider whether a phosphorylation site (Ser305Glu (Smith *et al*. 2023)) that we had previously characterized as underlying disease-strain specific seeding would have any deterministic role in seeding of hyperacetylated tau. Specifically, we showed that PL tau containing S305E can be seeded by PSP-tau but not by AD-tau seeds (Smith *et al*. 2023). To examine the interaction between acetylation and phosphorylation PTM with seeding propensity, we introduced the S305E substitution into PL+5K-Q tau (**Fig. 5a**). We first confirmed that the expression levels of the different tau variants, including the S305E containing PL+5K-Q, were equivalent to the other tau variants being tested. For each variant tested, the levels of soluble tau were similar (**Fig. 5b**). We observed complete ablation of seeding when the S305E mutation was combined with PL+5K-Q tau in the presence of AD-tau seeds (**Fig. 5c-d**; p<0.01), similar to the inhibition noted in parental PL tau carrying the S305E substitution (**Fig. 5c-d**; p<0.01). Interestingly, this new phospho-acetyl mimetic (PL+5K- Q/S305E) retained seeding responsiveness to PSP-tau seeds (**Fig. 5c-d**). This is consistent with our earlier data that parent PL tau with the S305E substitution can be seeded by PSP-tau but was generally not responsive to AD-tau seeds (Smith *et al*. 2023). None of the constructs showed any insoluble tau when left unseeded. These findings indicate hyperacetylation mimetic mutations did not alter the inherent prion specificity of Ser305Glu pseudo-phosphorylated tau for PSP-tau seeds. We next examined the binding of these variants to polymerized microtubules. In the presence of paclitaxel-stabilized microtubules, the levels of co-sedimented PL+5K-Q/S305E tau was considerably lower than P301L tau (**Fig. 5e-f**, p<0.0001 with paclitaxel; **Fig. 5g-h**, p=0.0014 without paclitaxel), though modestly higher than PL+5K-Q tau, indicating that both pseudo-acetylated and pseudo-phosphorylated tau associate less effectively to microtubules relative to PL-tau.

**Figure 5:**
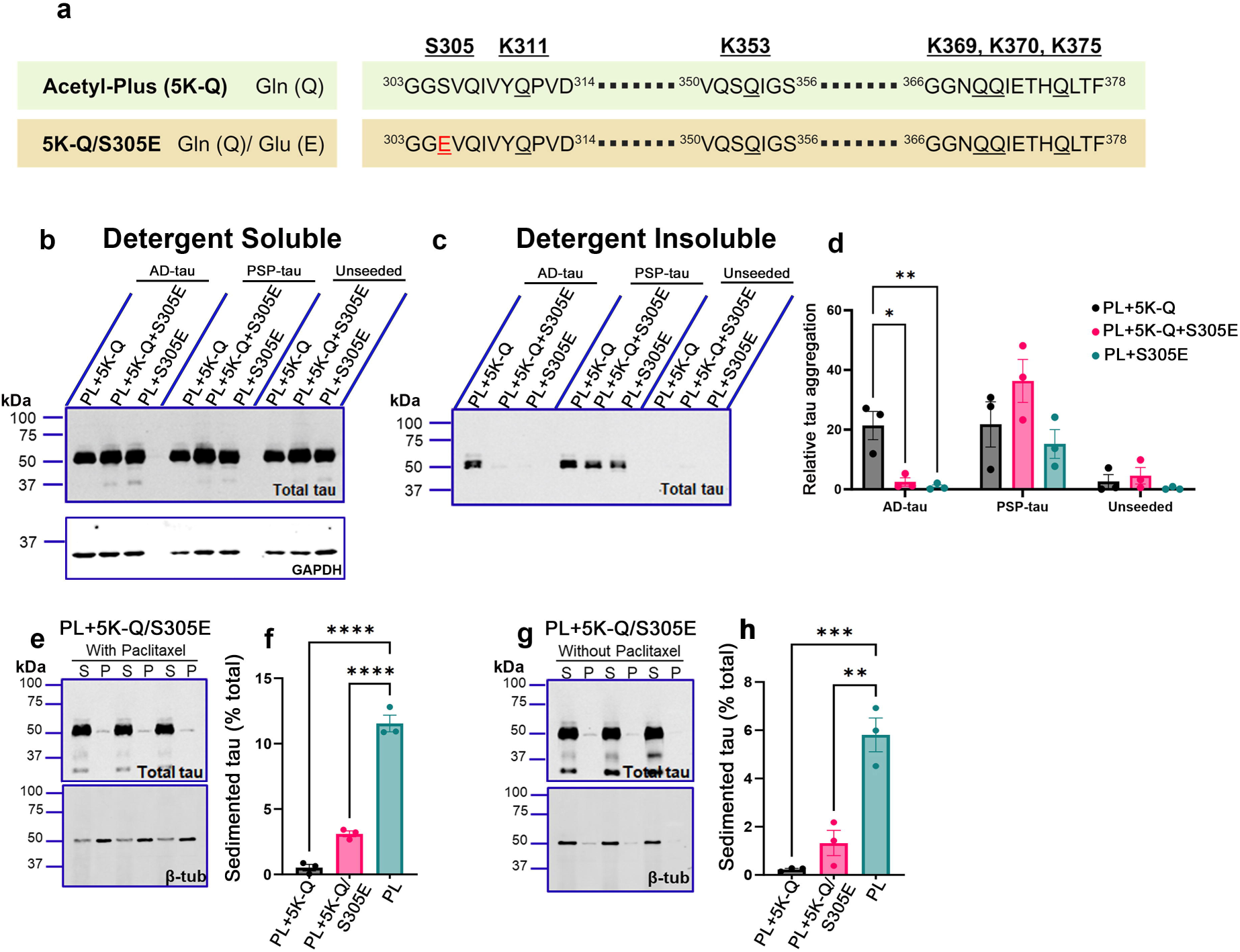
Presence of phospho-mimicking Ser305Glu modifies the ability of pseudo-acetyl 5K-Q tau to be differentially seeded by AD-tau and PSP-tau seeds. a. Linearized depiction of pseudo-acetyl PL+5K-Q tau carrying the phosphorylation-mimetic substitution on Ser305 (Ser➔Glu) generated on human 0N/4R P301L mutant tau. b-d. Seeded HEK293T cells expressing PL+5K-Q, PL+5K-Q+S305E and PL+S305E were fractionated into detergent-soluble and detergent-insoluble lysates and probed for total tau. Representative immunoblots (b, c) and relative tau aggregation (d) are shown. GAPDH immunoblot indicates protein loading control in the detergent soluble fraction. e-h. Representative blots and quantification of sedimentation of PL+5K-Q+S305E mimetic compared to PL+5K-Q and parent PL tau in the presence (e-f) and absence of paclitaxel (g-h). Blots were probed for total tau or β-tubulin (β-tub). All graphs represent mean ± S.E.M. N=3 for each experimental replicate. Graphical data for PL and PL+5K-Q tau shown in panels f, h correspond to data from Fig. 4b, d. Relative molecular masses (kDa) are indicated on the left of each blot. 2-way ANOVA with Dunnett’s multiple comparisons test, with single pooled variance. *p<0.05, **p<0.01, ***p<0.001, ****p<0.0001. PL, P301L tau; AD, Alzheimer’s disease; PSP, Progressive supranuclear Palsy; S, supernatant fraction; P, pellet fraction.

### 3.6. Analyzing pseudo-acetylated tau structure using *in silico* prediction

To try and gain more understanding of how acetylation could impact susceptibility to tau seed propagation, we performed in silico modeling analysis on the known filament core domains from AD and PSP (Fitzpatrick *et al*. 2017; Shi *et al*. 2021). We substituted the lysine residues at K311, K353, K369, K370 and K375 with Q residues to predict how the K→Q substitutions could potentially influence intramolecular interactions on tau compared to corresponding non-mutated (or WT) residues (**Fig. 6**). In AD-tau, the Q substitutions were predicted to cause subtle changes in the positioning of the amino acid side chains and affect interactions with neighboring residues, which is reflected in altered inter-molecular distances in K311Q and K370Q (**Fig. 6a, d**). Notably, there is a potential loss of a salt bridge interaction between K370 and D314, which could impact the folding (**Fig. 6d**). The positively charged and polar Lys370 was predicted to have strong interactions with surrounding polar, negatively charged Glu372 and Asp314 residues along with the uncharged Ser316. However, the uncharged and polar Lys370Gln residue will presumably incur weakened interactions because the positive charge that was associating with the negatively charged Glu372 and Asp314 is no longer there, potentially causing increased repulsion (2.4-3.4Å➔5.6Å) (**Fig. 6d**). For K353 within the AD-core, the positively charged and polar Lys353 residues has a strong interaction with the negatively charged, polar Asp358 residue. However, the uncharged and polar Lys353Gln residue potentially leads to a decrease in interaction between the negatively charged, polar Asp358 residue due to loss of charge (4.5Å➔5Å) (**Fig. 6b**). In some cases, contact with phospho-epitopes were also modified, which could potentially affect the availability of these Ser/Thr residues to PTM on AD-tau. For example, K353Q moves closer to Ser356 (6.0Å➔4.3Å) and K375Q moves away from Thr373 (3.8Å➔5.5Å) (**Fig. 6b, e**).

**Figure 6:**
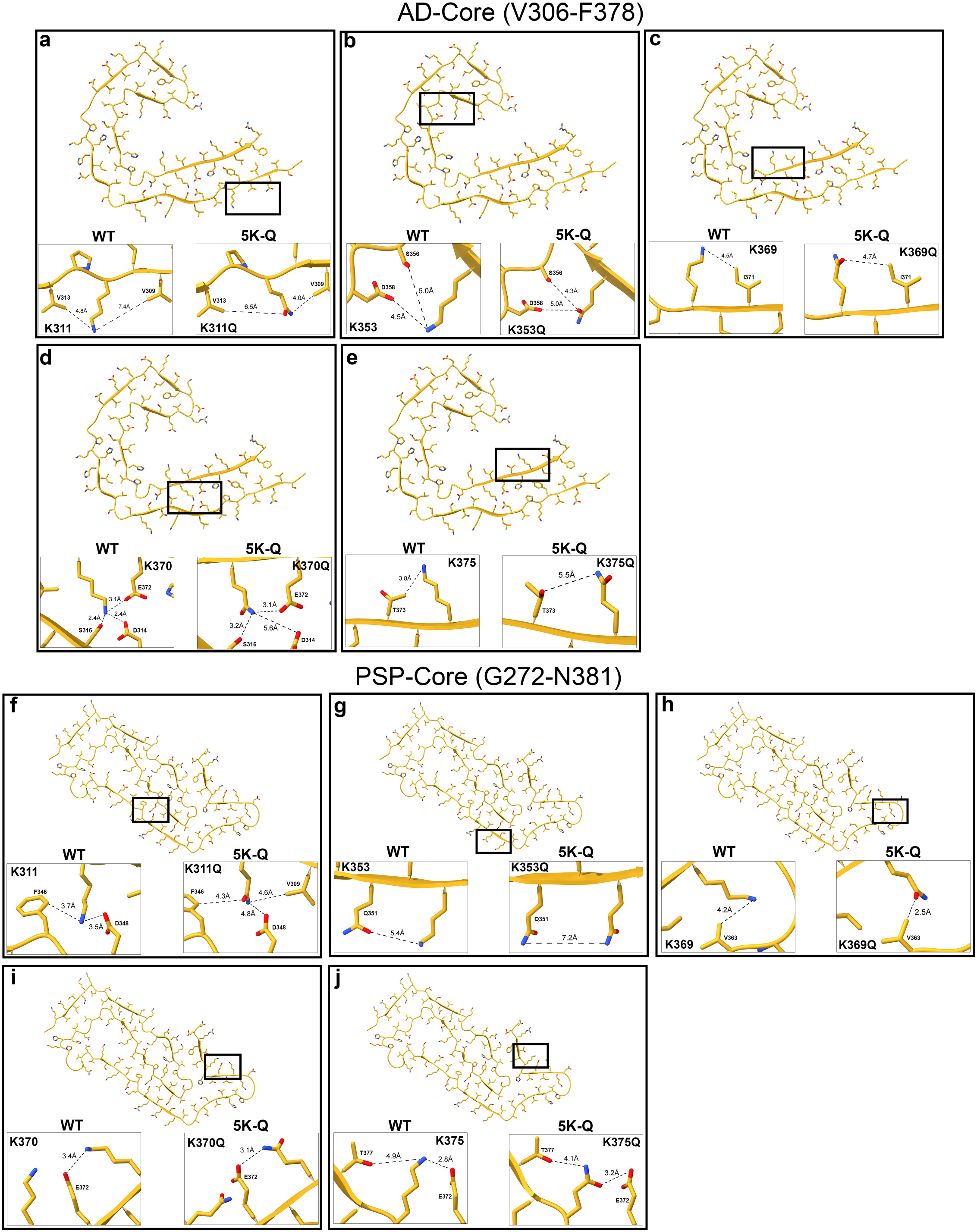
In silico predictive modeling of acetyl-mimetic tau contained in the tau core domains found in AD and PSP. In silico predictions of 5K-Q substitution or no substitution (WT) generated on the AD-tau core (a-e) (7QL4; amino acids V306-F378; (Lovestam *et al*. 2022)) and PSP-tau core (f-j) (7P65; amino acids G272-N381; (Shi *et al*. 2021)) using ChimeraX. AD, Alzheimer’s disease; PSP, Progressive supranuclear Palsy; WT, wild type; 5K-Q, Gln (Q) substitution on K311, K353, K369, K370, and K375 sites.

Comparing the corresponding pseudo-acetyl residues on the PSP-tau PDB framework (**Fig. 6f-j**), we found similarly that the K→Q substitutions showed subtle changes in interactions between neighboring residues, especially K311Q, K353Q and K369Q (**Fig. 6f-h**). For K311 within the PSP-core, the positively charged and polar Lys311 have strong interactions between the polar and negatively charged Asp348. However, the uncharged and polar Lys311Gln residue lost the charged interaction with Asp348, therefore potentially increasing the repulsion of the residues (2.8-3.5Å➔4.8Å) (**Fig. 6f**). Additionally, the positively charged and polar Lys369 likely does not interact well with the uncharged, non-polar Val363. At low pH, the repulsion likely increases due to the protonation of Val363. However, the uncharged and polar Lys369Gln residue lacks a charge, likely decreasing the repulsion with Val363 (3.5-4.2Å➔2.5Å) (**Fig. 6h**). In the in-silico models of contact prediction involving pseudo-acetylated PSP-tau, we did not observe any potential altered interactions with Ser/Thr residues. Although the individual changes in structure at each site are subtle, the combined impact may be more destabilizing to the filamentous tau aggregates.

## 4. Discussion

Specific acetylation patterns on tau are associated with disease stage-specific tau aggregation in tauopathies and thus are broadly thought to contribute to the progressive neurodegenerative cascade in tauopathies (Wesseling *et al*. 2020; Arakhamia *et al*. 2021). Previous research has demonstrated the pathological role of tau acetylation in the proline-rich region and MTBR R1-R2 domains (Min *et al*. 2010; Cook *et al*. 2014a; Min *et al*. 2015; Tseng *et al*. 2021; Shin *et al*. 2021; Chakraborty *et al*. 2023). Recently, specific acetylation events residing on MTBR R3-R4 domains spanning the tau filament core domain (K311, K353, K369, K370 and K375) were identified at high frequency in AD patients (Wesseling *et al*. 2020), leading us to examine whether presence of acetylation on these sites influenced templated tau aggregation. Given that the frequency of acetylation on these sites in the PSP-tau core is predicted to be lower from recent cryo-EM rendering, our secondary aim was to examine if acetylation on these sites led to strain-specific seeding of tau. Utilizing an experimental approach that we had previously used to examine the influence of phosphorylation PTM on tau seeding, we show that: (1) combinatorial acetyl-mimetic or acetyl-nullifying mutations were efficiently templated by both AD and PSP brain-derived tau seeds; (2) a combinatorial acetyl-null construct with alanine substitution spontaneously produces amorphous aggregates that are capable of seeding naïve PL tau template, and (3) strain specific features of the S305E phospho-variant dominated over hyperacetylated tau in producing a strain-like pattern when PL+5K-Q tau with S305E was exposed to AD-tau and PSP-tau seeds. Thus, our results demonstrate that phosphorylation mimetic mutation on S305 strongly influences the strain characteristics during seeding whereas, acetylation within the tau filament core, whether singly or combinatorially, has little impact on strain specificity. Overall, our data indicates that acetylation in the tau filament core domain is not a major driver in strain selection during tau seeding in the HEK293T cell model.

Many studies have established the pathogenic properties of acetylated tau in the context of tauopathies. Of the >30 potential AcK epitopes, several studies have shown that AcK174, AcK274, AcK280 and AcK281 are found in NFT from patient brains and underlie toxicity in mice (Irwin *et al*. 2012; Cohen *et al*. 2011; Tracy *et al*. 2016; Min *et al*. 2015). These AcK sites are located in the proline rich region and MTBR R1-R2, regions that are outside of the canonical tau filament core (Fitzpatrick *et al*. 2017). In a recent study using 4R tau, Cohen and colleagues demonstrated that PL tau protein that has an acetyl-nullifying K280→R substitution showed delayed aggregation kinetics whereas the pseudo-acetyl K280→Q showed modestly increased aggregation (Tseng *et al*. 2021). Another recent study showed that acetylation of 4R tau strongly inhibits in vitro fibrillization kinetics (Chakraborty *et al*. 2023), consistent with our cellular seeding data using the five combinatorial pseudo-AcK sites around MTBR R4. Notably, cooperative hyperacetylation on the KXGS motifs spanning the MTBR regions, specifically, K259/290/321/353Q substitution inhibited tau aggregation (Cook *et al*. 2014a), though individually K353Q did not influence tau fibrillization in vitro (Carlomagno *et al*. 2017). In addition, single site substitutions on K311 and K369 also did not substantially alter tau fibrillization (Carlomagno *et al*. 2017), suggesting that inter-AcK cooperativity is critical. Altogether tau acetylation has been proposed to underlie many cellular deficiencies, such as disruption of tau-microtubule interaction, impaired turnover and clearance, synaptic dysregulation and increasing the rate of tau fibrillization (Cohen *et al*. 2011; Min *et al*. 2010; Tracy *et al*. 2016; Haj-Yahya & Lashuel 2018). Notably, all of these studies have primarily assessed the spontaneous aggregation of tau. However, it still remains unknown if acetylation would be an early event underlying strain selectivity of tau or would be a later event occurring after tau has been misfolded into NFT. Our study is based on selected AcK sites that have been identified within the filament core domain of tau from AD patients using sensitive mass spectrometry techniques (Wesseling *et al*. 2020). Based on this information, we have asked the question whether acetylation of tau within the canonical tau filament core influences tau seeding when exposed to insoluble tau seeds derived from AD and PSP donors. Examination of single site mutations for each of these acetylation sites did not reveal any obvious changes in seeded aggregation when compared to parent PL tau. We also show that cooperative hyperacetylation on specific residues within the filament core domain (K311, K353, K369, K370 and K375) does not contribute to strain-specific tau seeding. Our data is consistent with the theory that acetylation is a late event that is downstream from seeding and spreading, reminiscent of the observations in post-mortem AD cases which reported that acetylation on these 5 sites is predominantly a late event in the tauopathy cascade, possibly occurring at or beyond Braak stage 5 (Wesseling *et al*. 2020). Given that seeding and templating has been thought to occur in earlier Braak stages leading to spread of tauopathy (Manca *et al*. 2023; DeVos *et al*. 2018), we speculate that acetylation on these 5 sites are not typically associated with triggering the initial seeding phase but would be associated more with end-stage conformational changes when the filaments are getting compacted into β-sheet structures later in the disease process.

Biochemical and neuropathologic data suggests that the predominant form of misfolded tau that is found in different tauopathies have unique structures (Chung *et al*. 2021). This finding has led to the strain hypothesis of tau, whereby tau adopts specific conformations that determines how it is templated and propagated along specific pathways and affecting specific cell types. This strain hypothesis could explain the intra-individual phenotypic, neuropathological and clinical diversity within each tauopathy as well as between different tauopathies (Vaquer-Alicea *et al*. 2021). Indeed, rodent modeling studies (Kumar *et al*. 2024; Dujardin *et al*. 2020; Kaufman *et al*. 2016; Holmes & Diamond 2014; Narasimhan *et al*. 2017) are now supported by elegant molecular-level cryo-EM data (Shi *et al*. 2021; Scheres *et al*. 2020) which show that native tau transforms into specific filamentous patterns that are unique to the clinical delineation of the tauopathy. While cellular and molecular studies establish how phosphorylation determines the seeding propensity of tau (Dujardin *et al*. 2020; Smith *et al*. 2023), less is known about acetylation as a determinant factor. Recent data shows that acetylation on K298/K294 (MTBR-R2 domain) and K311 (MTBR-R3 domain) could result in differential disease-specific tau strains based on tau isoforms (Trzeciakiewicz *et al*. 2020; Chakraborty *et al*. 2023). Specifically, acetylation on K298/294 delays 4R tau aggregation whereas AcK311 favors 3R tau aggregation. Additionally, K280Q substitution on P301S mutant tau abrogates its natural propensity to seed and form inclusions in cellular assays (Ajit *et al*. 2019). Our data now provides additional evidence that cooperative acetylation of AD-associated AcK sites within the tau filament core does not majorly influence seeded tau aggregation.

Proteomic and neuropathologic data from human patient samples suggest that cooperativity between different PTM sites determine tau misfolding (Wesseling *et al*. 2020; Kyalu Ngoie Zola *et al*. 2023). Acetylation mediated downstream activity (such as those occurring in histones) generally follow phosphorylation signals, which act as initiating cellular signaling stimuli in normal homeostatic conditions in the nucleus. Whether such a directional modality exists in the life cycle of tau as posited by Wesseling and colleagues (Wesseling *et al*. 2020) is unknown. In addition, Lysine residues are modified not only by acetylation, but also by methylation, ubiquitination, sumoylation and neddylation in a mutually exclusive manner, thus increasing the complexity of the PTM code in the lifecycle of tau. Additionally, a mouse model expressing the P301S tau with K280→Q substitution was unexpectedly protected from tau-induced early mortality and displayed striking levels of attenuated phosphorylation on tauopathy-associated epitopes (Ajit *et al*. 2019), suggesting that the presence of P301S (a potentially phosphorylating mutation) could alter how acetylation mimetics regulate tau aggregation. Thus, with regard to tau, this is a complex relationship as specific PTM could alter tau aggregation in both positive and negative manner and that one type of PTM could preclude another type, thus complicating the outcome (Alquezar *et al*. 2020). Some molecular evidence shows such PTM-site cooperativity in the context of tau. For example, pseudo-acetyl K259/290/321/353Q tau reduces tau fibrillization in vitro and further modifies the phosphorylation on specific Ser epitopes in its vicinity (Cook *et al*. 2014a; Carlomagno *et al*. 2017), suggesting cooperativity between phosphorylation and acetylation status. We provide *in silico* modeling data to suggest that hyperacetylation in the AD-tau filament core could potentially affect phosphorylation on neighboring Ser/Thr residues. Our previous study and the current work also demonstrate that the phosphorylation mimicking mutation Ser305Glu strongly determines susceptibility to AD-tau versus PSP-tau seeds. We find that this determinant factor is not modified by pseudo-acetyl motifs, and that pseudo-acetylation did not itself modify strain susceptibility. Moreover, PL tau variants incapable of acetylation retained high susceptibility to phosphorylation at the AT8 site. Overall, our data suggest that phosphorylation on specific epitopes has a stronger impact on strain-like characteristics of tau propagation than acetylation.

One of the most abundant amino acids in the AD filament core is lysine (14.3%), underscoring the importance of lysine site modifications in disease pathogenesis. *In silico* data presented indicates that for the AD filament core, the two sites that emerged as of particular interest due to the increases in interacting residues and location were K353 and K370. On the other hand, altering acetylation potentially affected K311 and K369 more strongly than the other residues on the PSP-tau filament core. Substitution of Lys with Alanine in 5K-A would reduce the net positive charge, possibly leading to its reduced solubility. Indeed, cryo-EM studies suggest that neutralization of the positive charges of the lysines within the cross-β helix is necessary to form stable filaments (Zhang *et al*. 2019). Tau filaments from different tauopathies confirm that neutralizing the repulsion of positively charged lysine residues through acetylation facilitates formation of parallel in-register stacking of β-strands (Arakhamia *et al*. 2020). It is also possible that the 5K-A substitution reduces tau stability by disrupting Lys-related hydrophobic interactions or by altering flexibility afforded by presence of Lysines in loop structures. Several of the Lysine residues on tau can form hydrogen bonds and salt bridges to stabilize the core filament (Fitzpatrick *et al*. 2017). Additionally, cryo-EM ‘densities’ around lysine residues (e.g., K311 on AD filament) could be due to mono- or poly-ubiquitinated chains, resulting in stable β-strand stacking along the long axis of tau fibrils (Arakhamia *et al*. 2020). In PSP, a salt bridge formed by K311 as well as prominent densities on solvent-exposed K280/281 are critical to stabilize the tau filament (Shi *et al*. 2021). An interesting aspect is that while the selective patterns of Lys acetylation in the filament core is distinctive between AD-tau and PSP-tau, with AD-tau displaying higher acetyl occupancy along the filament core (Kametani *et al*. 2020; Arakhamia *et al*. 2020), the combinatorial and individual acetyl mimics used in this study did not necessarily show strain-specific differences in seeding. For example, the K311 side chain is buried within PSP-tau, while in AD-tau, it is solvent-accessible as it points away from filament core and is frequently acetylated. We had expected that such strain-specific acetylation patterns would form a PTM code that would determine tau templating. Based on our experimental data here, we conclude that acetylation is likely a secondary modifier of tau filament structure and additional PTMs, such as phosphorylation, play a primary deterministic role in the templating process.

Together with our earlier report (Smith *et al*. 2023), data presented here indicates that phosphorylation mimetic on the Ser305 position is associated with imparting strain-like characteristics during tau seeding, even in the presence of the combinatorial acetyl-modified epitopes. This specific residue is intriguing as its nucleotide sequence also determines the splicing of tau, and specific Ser305 mutations found in human tauopathy patients favor 4R tau (Rossi & Tagliavini 2015). In our seeding data, Ser305Glu containing acetyl-site modified tau was templated by the 4R tauopathy-derived PSP-tau seeds but not by 3R/4R mixed tauopathy-derived AD-tau seeds. This is consistent with data that AD cases did not seed tau biosensor cells carrying the tauopathy-driving Ser305Asn mutation (Watamura *et al*. 2025). On the other hand, brain extract from a human Ser305Asn mutation carrier seeded the Ser305Asn biosensor cells, confirming that mutations on Ser305 impart strain specificity. In iPSC models, the Ser305Asn mutation causes neuronal hyperexcitability, synaptotoxicity and astrocyte activation, as well as increased synaptosomal accumulation of hyperphosphorylated tau (Bowles *et al*. 2024). While Ser305 mutations to Asn and Ile have been confirmed to alter the 3R:4R tau ratio, the role of these mutations in ablating Ser305 phosphorylation and determining strain specificity during seeding requires further work.

A biochemically and physiologically relevant issue is distinguishing our current work using acetyl-mimetics with naturally phosphorylated ac-K tau. Most of the work cited here, as with ours, have used genetically modified mutated constructs to prevent (K→R or A) or mimic (K→Q) Lys acetylation on specific epitopes. However, there are studies that have used biochemically modified tau to ascertain how acK-tau influences tau seeding and aggregation. For example, a study used acetyl coA-modified recombinant P301L tau seeds in cell culture, finding that acetylated seeds did not influence aggregation propensity (Tseng *et al*. 2021). On the other hand, co-expressing the acetyltransferase Creb-binding protein/p300 to acetylate both the seed and the template increased seeded tau aggregation in cells. Additionally, pre-acetylating K267, K274 and K280 in a modified K18 peptide using N-succinimidyl acetate increased seeding of this aggregated peptide in a FRET-based assay (Li *et al*. 2023). The main difference between naturally acetylated tau is that the percentage of tau that is modified at any one site will be variable and the modification is dynamic. This complexity cannot be modeled by mimetic mutations.

One limitation of our study is that we used the FTD-associated PL tau to examine tau seeding. We have previously shown that WT tau is not receptive to templated aggregation in these cellular assays (Smith *et al*. 2023) which is supported by similar assays from other research groups using P301L or P301S backbone (Lathuiliere *et al*. 2023; Holmes & Diamond 2014). Another limitation is that our analysis is done using 0N/4R tau and given the newly emerging data on the differential toxicity of 3R and 4R tau in different tauopathies (Chakraborty *et al*. 2023; Trzeciakiewicz *et al*. 2020; Schoch *et al*. 2016), future studies will need to consider the relative seeding propensity of acetylated 3R and 4R isoform tau. Detailed mass spectrometry information on how acetylation of neighboring residues in the acetyl-modified P301L tau would give us further insights into self-templating properties of tau in disease. Lastly, we do not know whether the cell assay system we use faithfully propagates the AD and PSP strains of misfolded tau. The assay primarily reports seeding competency of the enriched insoluble tau fractions from AD and PSP brains and may not report faithful seed propagation.

In conclusion, our study shows that neither acetylation mimicking mutations nor acetylation-nullifying mutations in the filament core domain of PL tau modulate susceptibility to AD and PSP strains of tauopathy in forming tau aggregates in cellular seeding model. Our data suggest that pseudo-phosphorylation of PL tau at Ser305 is a strong modifier of susceptibility to seeding by AD and PSP-tau seeds irrespective of acetylation mimicking mutations within the tau filament core region, indicating that acetylation within the core filament could be secondary events in the lifecycle of tau in neurodegenerative diseases.

## Abbreviations

AcK: acetylated Lysine
AD: Alzheimer’s disease
AD-tau: detergent-insoluble tau filaments isolated from AD patients
PL+5K-Q: Gln (Q) substitution on 5 Lysine (K) sites in 0N/4R P301L tau
PL+5K-R: Arg (R) substitution on 5 Lysine (K) sites in 0N/4R P301L tau
PL+5K-A: Ala (A) substitution on 5 Lysine (K) sites in 0N/4R P301L tau
CBD: corticobasal degeneration
Immuno-EM: immunogold electron microscopy
MT: microtubules
MTBR: microtubule binding repeat domains
NFT: neurofibrillary tangle
PL: tau 0N/4R P301L tau
PSP: progressive supranuclear palsy
PSP-tau: detergent-insoluble tau filaments isolated from PSP patients
PTM: post-translational modification
p-tau: phosphorylated tau

## Author Contributions

PC: Conceptualization, Funding acquisition, Project administration, Supervision, Validation, Visualization, Writing—original draft, Writing—review & editing; DRB: Conceptualization, Writing—review & editing; EDS: Investigation, Data collection and analysis, Methodology, Writing—& editing; JT, QV: Investigation, Data Collection and analysis; SP, RM, MM: Resources, Writing—review. All authors have approved the manuscript and agree with its submission.

## Acknowledgements

We thank the University of Florida Neuromedicine Human Brain and Tissue Bank for sharing human brain samples (funded in part by NIA P30AG066506). S.P. is supported by the Charlotte and Howard Zimmerman rising star professorship at the Norman Fixel Institute for Neurological diseases

## Funding and Additional Information

The work was supported partially by NIA R01 AG078734 (PC, DRB). EDS was supported by the predoctoral T32NS082168. JT is supported by the predoctoral T32AG061892.

## Institutional Approval

The study was conducted according to the guidelines of the Declaration of Helsinki and approved by the Institutional Review Board of the University of Florida (IRB201600067). Use of recombinant DNA was approved by the University of Florida Environmental Health & Safety Office.

## Data Availability Statement

The data supporting the findings of this study are available on request from the corresponding author. AAV constructs will be available directly from Genscript on request made to the author and completion of institutional MTA.

