## Supplemental Tables for "Acetylation mimetic and null mutations within the filament core of P301L tau have varied effects on susceptibility to seeding and aggregation"

**Table S1. Demographics and diagnoses of the patient cohorts used in this study. Case IDs correspond to individually characterized brains in the University of Florida Neuromedicine Human Brain and Tissue Bank. Experimental Group denotes the designation for this study. NP Dx1, primary diagnosis based on neuropathology; NP Dx2 and NP Dx3, secondary diagnoses based on neuropathology: Thal phase, burden of immunostained amyloid deposits in cortical and subcortical area; Braak stage, and CERAD score, neuritic plaque frequency; APOE, Apolipoprotein E; AD, Alzheimer's disease; CAA, cerebral amyloid angiopathy; PSP, Progressive supranuclear Palsy. N/A: Not applicable; ND, not done; PMI: Post-mortem interval.**

| Experimental Group | Post-mortem Diagnosis | Case ID | NP Dx1 | NP Dx2 | NP Dx3 | Thal stage | Braak stage | CERAD score | APOE genotype | Sex | Age | PMI |
|---|---|---|---|---|---|---|---|---|---|---|---|---|
| AD-A | AD | A19-015 | AD high | CAA widespread, mild | N/A | 4 | V | frequent | 3/4 | m | 77 | 5 |
| AD-B | AD | A19-027 | AD high | CAA widespread, moderate | N/A | 5 | VI | frequent | 3/4 | m | 63 | 2 |
| PSP-A | PSP | A22-040 | PSP | AD low | CAA focal, mild to moderate | 3 | I | none | ND | m | 70 | 19.5 |
| PSP-B | PSP | A21-041 | PSP | AD intermediate | CAA widespread, severe | 5 | III | moderate | 4/4 | m | 72 | 27 |

**Table S2. Antibodies used in this study**

| Antibody name | Specificity | Host Species | Dilutions | Source |
|---|---|---|---|---|
| Recombinant anti-tau | Mouse and human tau | Rabbit monoclonal | 1:10000 (WB) | Abcam Cat# ab254256, RRID:AB_2894402 |
| CP27 | human tau (130-150) | Mouse monoclonal | 1:500 (EM) | Gift from Dr. Peter Davies (Duff et al., 2000; RRID:AB_2716722) |
| CP13 | pSer202 | Mouse monoclonal | 1:1000 (WB) | Gift from Dr. Peter Davies (Weaver et al., 2000; RRID:AB_2314223) |
| PHF1 | pSer396/pSer404 | Mouse monoclonal | 1:1000 (WB) | Gift from Dr. Peter Davies (Greenberg et al., 1992; RRID:AB_2313687) |
| AT8 | pSer202/pT205 | Mouse monoclonal | 1:1000 (WB) | Thermo Fisher Scientific Cat#MN1020, RRID:AB_223647 |
| AT180 | pThr231 | Mouse monoclonal | 1:1000 (WB) | Thermo Fisher Scientific Cat# MN1040, RRID:AB_223649 |
| AT270 | pThr181 | Mouse monoclonal | 1:1000 (WB) | Thermo Fisher Scientific Cat# MN1060, RRID:AB_223652 |
| AC-15 | Actin | Mouse monoclonal | 1:1000 (WB) | (Abcam Cat# ab6276, RRID:AB_2223210) |
| TUB 2.1 | β-tubulin | Mouse monoclonal | 1:1000 (WB) | (Sigma-Aldrich Cat# T4026, RRID:AB_477577) |
