## Supplemental Figures and Legends for "Acetylation mimetic and null mutations within the filament core of P301L tau have varied effects on susceptibility to seeding and aggregation"

**Supplementary Table Legends**.

**Supplementary Table S1**. Demographics and diagnoses of the AD and PSP brain donors.

Demographics and diagnoses of the patient cohorts used in this study. Case IDs correspond to individually characterized brains in the University of Florida Neuromedicine Human Brain and Tissue Bank. Experimental Group denotes the designation for this study. NP Dx1, primary diagnosis based on neuropathology; NP Dx2 and NP Dx3, secondary diagnoses based on neuropathology: Thal phase, burden of immunostained amyloid deposits in cortical and subcortical area; Braak stage, and CERAD score, neuritic plaque frequency; APOE, Apolipoprotein E; AD, Alzheimer's disease; CAA, cerebral amyloid angiopathy; PSP, Progressive Supranuclear Palsy. N/A: Not applicable; ND, not done; PMI, post-mortem interval.

**Supplementary Table S2**. Antibodies used in this study.

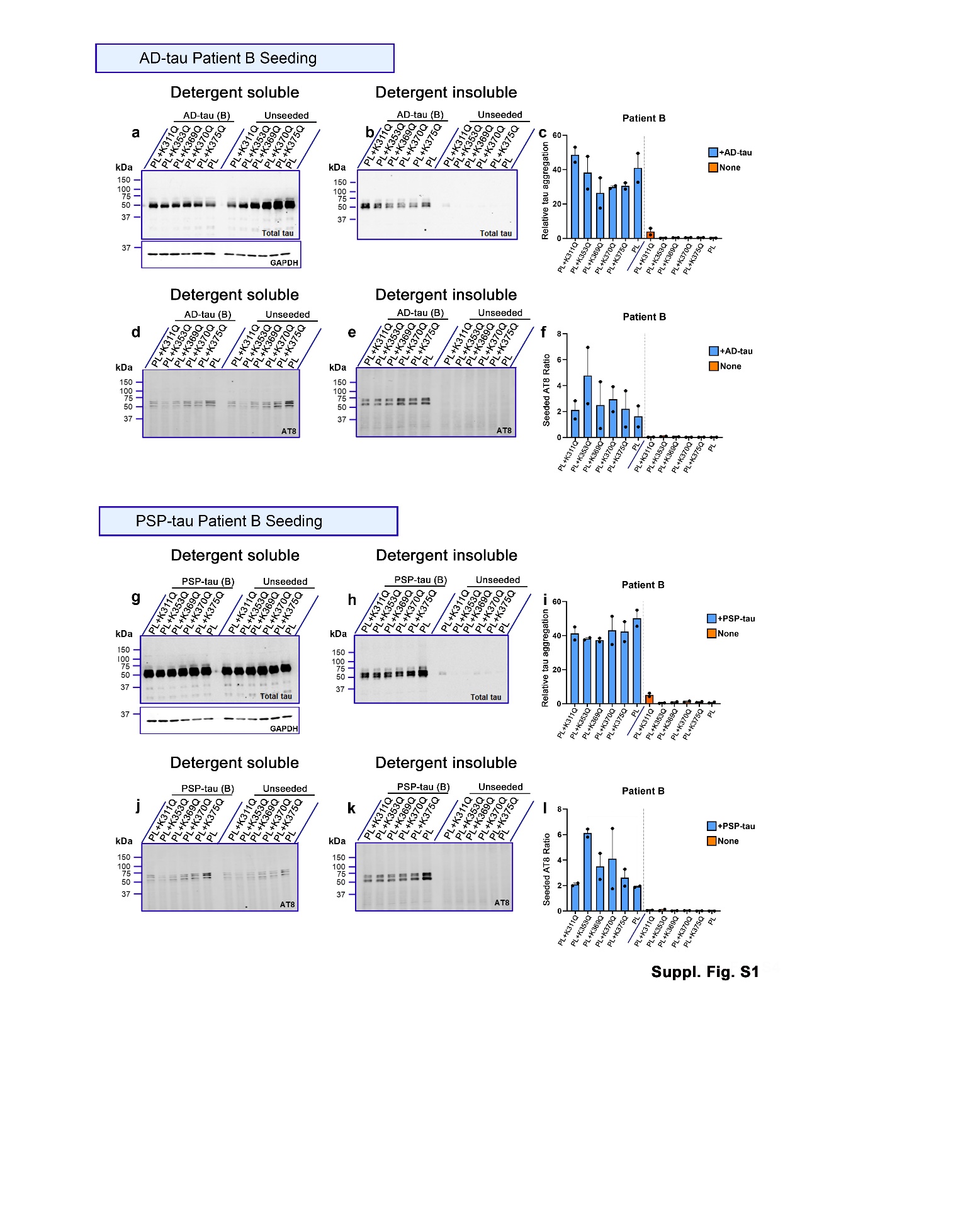

**Figure S1:** **Comparative profiles of** **individual pseudo-acetyl tau variants seeded with AD-tau and PSP-tau seeds.**

a-f. Representative immunoblots and quantitation depicting relative tau aggregation (a-c) and seeded AT8 ratio (d-f) of individual acetyl-mimetic tau seeded with AD:Patient B tau seeds. g-l. Representative immunoblots and quantitation depicting relative tau aggregation (g-i) and seeded AT8 ratio (j-l) of individual acetyl-mimetic tau seeded with PSP:Patient B tau seeds. Relative molecular masses (kDa) are indicated on the left of each blot. HEK293T cells expressing tau variants but not seeded shown as ‘Unseeded’ lanes and ‘None’ in graphs. N=2 for each experimental replicate.

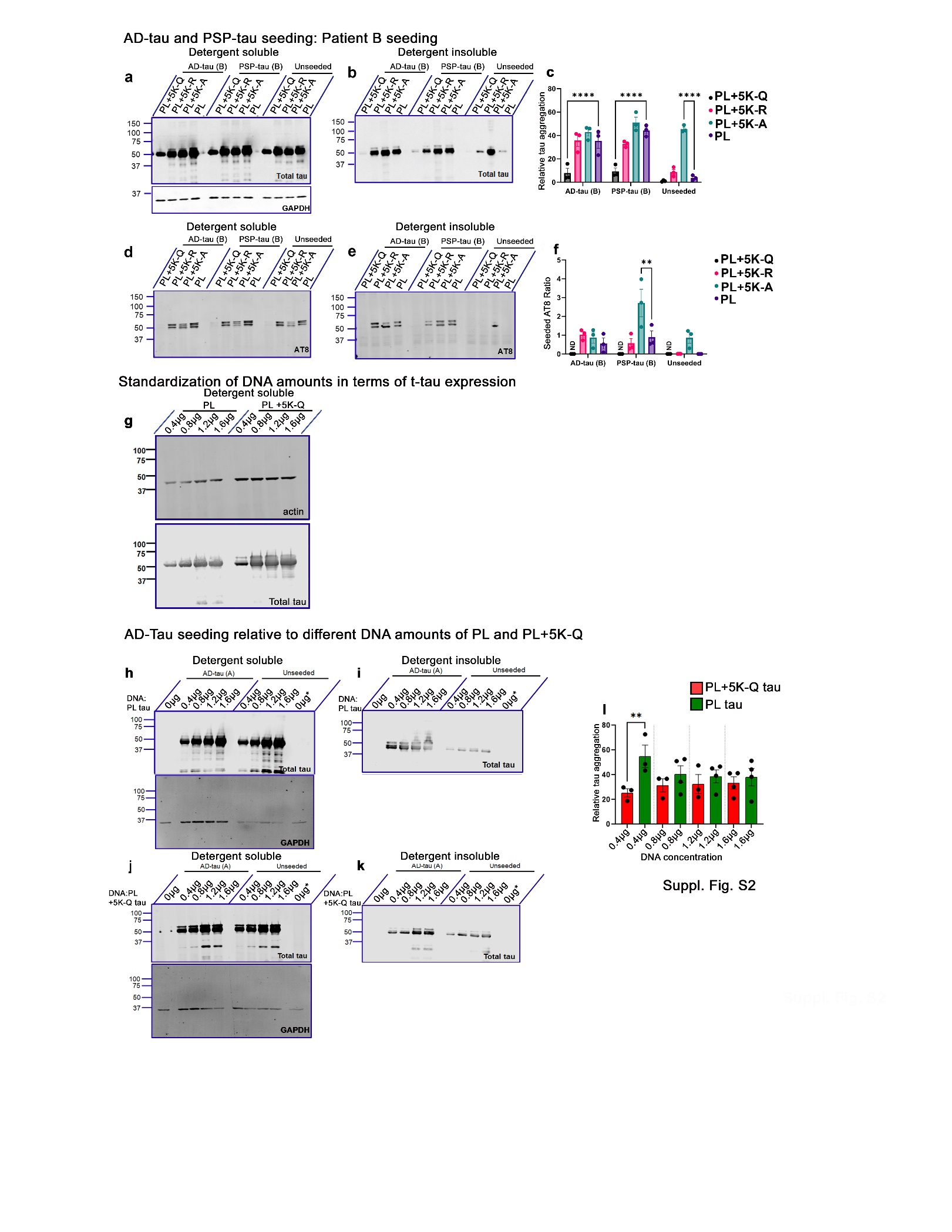

**Figure S2:** **Comparative effect of combinatorial acetyl variants in the tau core domain on seed-induced PL tau aggregation**.

a-f. HEK293T cell seeding assay using the acetyl variants seeded with AD-tau seeds or PSP-tau seeds. Samples were fractionated into detergent-soluble and detergent-insoluble lysates and probed for total tau and p-tau (AT8). Representative immunoblots (a, b, d, e), relative tau aggregation (c) and seeded AT8 ratio (f) of acetyl-tau variants seeded with AD:Patient B tau seeds or PSP:Patient B tau seeds shown. All graphs represent mean ± S.E.M. N=3 for each experimental replicate. GAPDH immunoblot indicates protein loading control. 2-way ANOVA with Dunnett’s multiple comparisons test, with single pooled variance. ****p<0.0001. g-l. Relative standardization of PL tau and PL+5K-Q tau expression levels by varying plasmid DNA used in transfection of HEK293T cells as indicated on top of lane (g). Following this, HEK293T cell seeding assay was carried out using varying amounts of PL tau DNA (h-i) or PL+5K-Q tau DNA (j-k) transfected and subsequently seeded with AD-tau seeds. Samples were fractionated into detergent-soluble and detergent-insoluble lysates and probed for total tau. Representative immunoblot (h-k) and relative tau aggregation (l) of acetyl-tau variants seeded with AD:Patient A tau seeds. Asterisk with 0µg denotes 1.6µg of GFP plasmid DNA. All graphs represent mean ± S.E.M. N=3-4 experimental replicates. GAPDH immunoblot indicates protein loading control. 1-way ANOVA with Tukey’s test. **p<0.01.

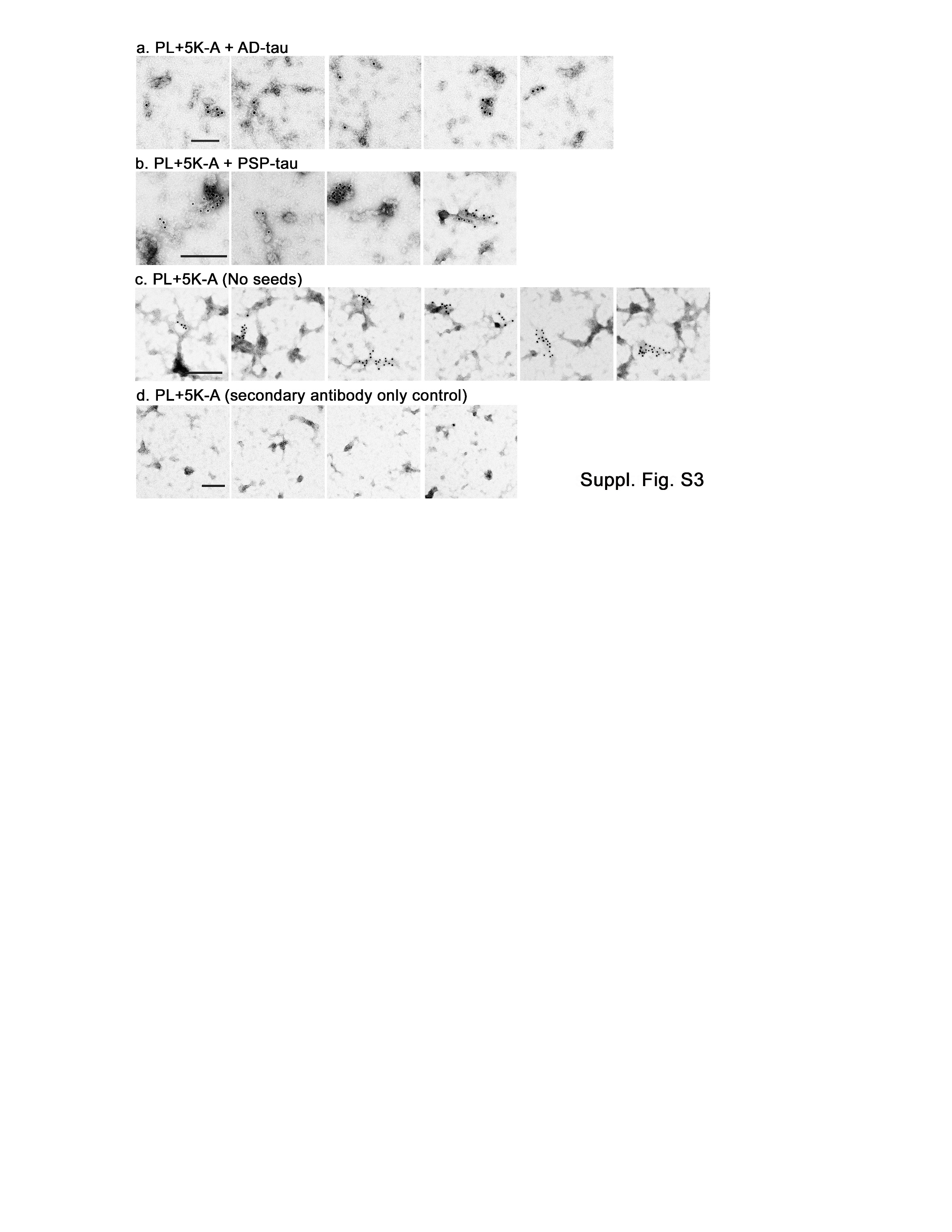
**Figure S3**: **Additional immuno-EM images for 5K-A tau aggregates**. a-c. Representative immuno-EM images of detergent-insoluble PL+5K-A tau aggregates from HEK293T cells that were seeded with AD-tau (a), PSP-tau (b) or left unseeded (c). d. Representative Immuno-EM images of 5K-A tau exposed to secondary antibody only. All preparations were stained with 10nm gold conjugated secondary antibody followed by negative staining with 1% uranyl acetate.

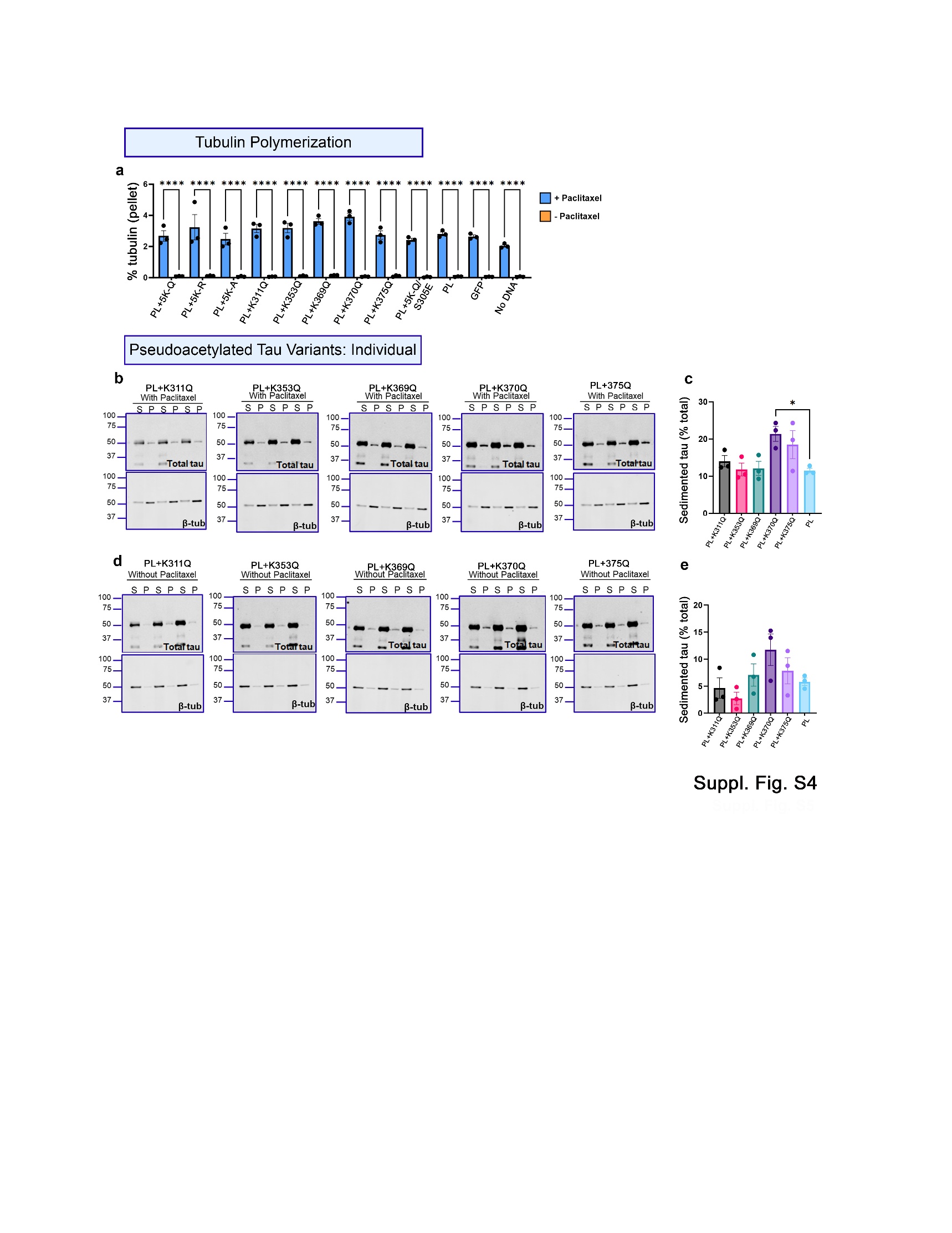

**Figure S4:** **Microtubule binding of individual acetyl-modified tau variants**.

HEK293T cells were transfected with individual acetyl-modified tau variants as indicated and cell lysates used for microtubule binding assay. a. Quantification of the tubulin polymerization efficiency in the presence of different acetyl tau variants with or without Paclitaxel. b-c. Representative blots and quantification of sedimented acetyl-modified tau variants compared to parent PL tau in the presence of paclitaxel. d-e. Representative blots and quantification of sedimented acetyl-modified tau variants compared to parent PL tau in the absence of paclitaxel. Blots were probed for total tau or β-tubulin (β-tub). 1-way ANOVA with Dunnett’s multiple comparisons test, with single pooled variance. **p<0.01, ***p<0.001, ****p<0.0001. S= supernatant fraction; P= pellet fraction. All graphs represent mean ± S.E.M. N=3 experimental replicates.
